# Temporal tuning of switch-like virulence expression resolves environmental uncertainty through phenotypic heterogeneity

**DOI:** 10.1101/2025.07.01.662634

**Authors:** Madison Spratt, Keara Lane

## Abstract

Gene regulatory networks often evolve in the face of environmental uncertainty, as stimuli are rarely precise and uniform enough to make all or nothing responses advantageous. Virulence gene regulation in intracellular bacterial pathogens is shaped by unique selective pressures to resolve this uncertainty, as host environments are dynamic, hostile, and heterogeneous. Here, we investigate the regulation of *Salmonella* Pathogenicity Island 2 (SPI-2), a set of genes required for intracellular replication of the enteric pathogen *Salmonella* Typhimurium (STm), to evaluate whether it has evolved the capacity to tune transcriptional responses to intracellular-like environments. Using live-cell reporters and smFISH in *in vitro* inducing environments, we identify bimodal expression of SPI-2 in clonal populations. We show cells progressively transition into a SPI-2 expressing state, with multi-generational variation in the timing at which this transition occurs. The rate of these single-cell transitions are modulated by the strength of intracellular-like signals and environmental manipulations of growth rate. We determine that heterogeneity originates at the level of sensor kinase SsrA signaling and is dependent on response regulator SsrB concentration. We identify switch-like dynamics in SPI-2 activation and link single-cell responses to features of cell growth and lineage. Combined, our results show that the fraction of cells expressing SPI-2 in a given timeframe is probabilistically tuned to the likelihood they are experiencing an intracellular environment, a strategy that resolves uncertainty in ambiguous environments.

## INTRODUCTION

Bacteria must survive and proliferate in dynamic and diverse conditions, requiring accurate detection of environmental change to mount appropriate functional responses. However, these responses must occur as bacteria navigate environments across a range of strengths, complexities, and stabilities; often forcing bacteria to make critical regulatory decisions with suboptimal environmental information (Bakshi et al., 2021; Mattingly et al., 2021; Nguyen et al., 2021; Spratt & Lane, 2022). Despite this frequent ambiguity, bacterial gene regulatory mechanisms manage to effectively avoid both aberrant induction and failed responses, an evolutionary feat that is particularly impressive when further considering the time constraints imposed by the life-or-death nature of these responses (Mitosch et al., 2017; Schultz et al., 2017; Uphoff, 2018). Here, we investigate the strategies a bacterial pathogen that routinely experiences environmental uncertainty has evolved to promote robust responses and ensure bacterial survival and proliferation.

The capacity to tune functional responses to environmental inputs is likely required to resolve this environmental uncertainty. Tuning has been identified in a wide variety of biological systems, including in bacterial stress responses (Locke et al., 2011; Lagage et al., 2023) and chemotaxis (Jasuja et al., 1999; J. P. Moore et al., 2024). One level at which this may occur is in the fraction of individuals that will respond, creating functionally distinct subpopulations in an isogenic population. The formation of such subpopulations is broadly referred to as phenotypic heterogeneity, resulting from the amplification of stochastic noise in gene expression (Elowitz et al., 2002; Raj & van Oudenaarden, 2008). Phenotypic heterogeneity is widely observed in bacterial species, allowing for division of labor or bet-hedging based strategies that promote population-level survival (Ackermann, 2015). Tuning may also occur in temporal features of the response of individual bacteria (Lagage & Uphoff, 2020; Levine et al., 2013; Locke et al., 2011; Patange et al., 2018; Sampaio et al., 2022), requiring quantification of the response in single cells as they transition into new environments. While many bacterial response systems have been extensively characterized at the population level, investigating these systems through a dynamic, single-cell lens is often necessary to reveal what features of the response are tunable. Doing so can provide mechanistic insight into gene regulation and lead to hypotheses about the evolutionary pressures that shaped the regulatory network, as has been shown in model systems such as the *E. coli* lac operon (Ozbudak et al., 2004; Kaiser et al., 2018; Julou et al., 2020) and the *B. subtilis* competence response (Maamar et al., 2007; Süel et al., 2007).

Complex environmental transitions are a hallmark of the lifecycle of bacterial pathogens, offering unique selective pressures that necessitate the evolution of tunable regulatory strategies. To survive, replicate, and disseminate within a host, bacteria express virulence genes encoding features such as secretion systems, flagella, and toxins (Deng et al., 2017; Diard & Hardt, 2017; Wu et al., 2008). Expression of these features are necessary for effective colonization and growth within their hosts, but misexpression carries significant costs, such as energetic deficits and host detection (Davis, 2020). Pathogens routinely traverse diverse environments as they move through a host, but typically require activation of particular virulence genes only within a specific niche. Single-cell approaches have revealed some strategies pathogens use to balance this necessity and cost tradeoff, particularly in systems that regulate invasion of host cells in enteric pathogens. For example, the invasion regulating thermosensor RovA in *Yersinia* (Nuss et al., 2016) and flagella regulation (Koirala et al., 2014) in *Salmonella enterica* serovar Typhimurium (STm) are bistable regulatory systems that tune the fraction of virulence expressing cells to stimulus strength. A similar strategy is used to a different effect in the regulation of STm SPI-1, a virulence system required for epithelial cell invasion in which bistable expression allows for a division of labor strategy that promotes gut lumen colonization while offsetting the growth costs of SPI-1 expression (Ackermann et al., 2008; Diard et al., 2013; Sturm et al., 2011).

Following uptake into a host cell, intracellular bacterial pathogens encounter a dynamic and hostile environment within the phagosome. Here, newly isolated bacteria must rapidly adapt to increasing acidity and nutrient deprivation as the host cell attempts to neutralize them (H.J. Lee et al., 2020; Sivaloganathan & Brynildsen, 2021). The pathogen STm has evolved to replicate intracellularly through activation of a set of genes known as Salmonella Pathogenicity Island 2 (SPI-2) (Cirillo et al., 1998; Han et al., 2024). SPI-2 encodes a Type 3 Secretion System (T3SS) and a set of effector proteins, which are secreted into the host cell cytosol to promote maintenance and maturation of the phagosome into a *Salmonella* Containing Vacuole (SCV) and intracellular replication (Figure 1A) (Jennings et al., 2017). A large body of research has elucidated the SPI-2 gene regulatory network as well as the primary environmental signals that activate it within the host cell. SPI-2 encompasses 32 genes within the island itself, encoded within six operons (Ochman et al., 1996; Shea et al., 1996; Tomljenovic-Berube et al., 2010) (Figure 1A). The genes that encode the structural and functional components of SPI-2 are regulated by SsrB, a response regulator in the two-component system (TCS) SsrAB (Garmendia et al., 2003). SsrAB activation is primarily controlled by the activation of two other TCSs, PhoPQ and OmpR/EnvZ (A. K. Lee et al., 2000; Bijlsma & Groisman, 2005; Fass & Groisman, 2009), that sense environmental signals, including reductions in pH and magnesium limitation (Deiwick et al., 1999; Nikolaus et al., 2001; Rappl et al., 2003; Löber et al., 2006). These then activate transcription and translation of *ssrAB*, which also contains a putative autoregulatory binding site (Tomljenovic-Berube et al., 2010). SsrB binds the promoters of all downstream SPI-2 genes and promotes transcription via RNA polymerase recruitment and displacement of the global negative regulator H-NS (Tomljenovic-Berube et al., 2010; Walthers et al., 2011). Multiple negative regulators target SsrB: the iron-sensing transcription factor Fur negatively regulates SsrB at the transcriptional level (E. Choi et al., 2014), while the EllaNTR protein, encoded by *ptsN*, directly interacts with SsrB and is proposed to bind and sequester SsrB and PhoP to prevent premature downstream gene expression (J. Choi et al., 2010, 2019).

**Figure 1.**
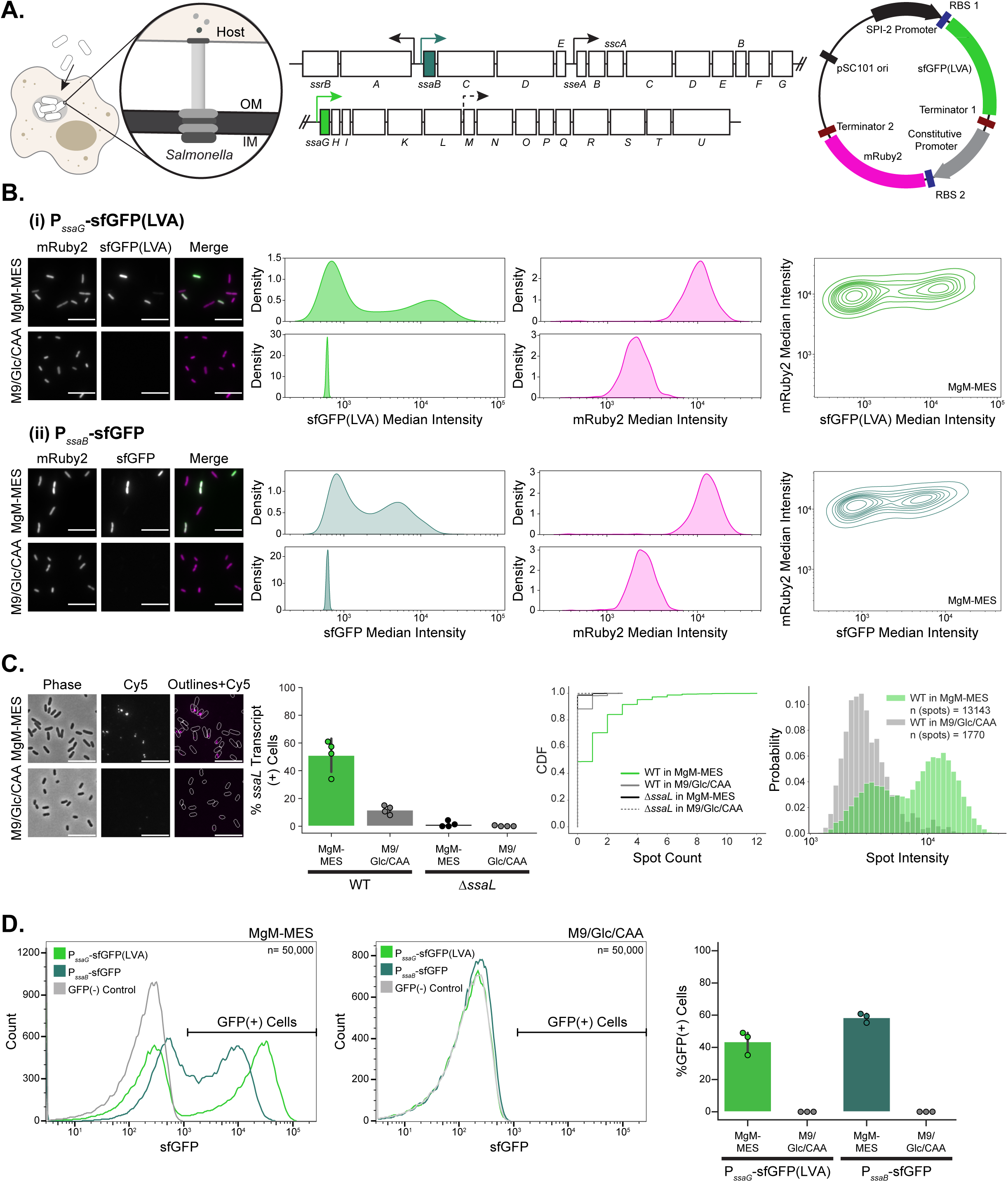
*Salmonella* SPI-2 is bimodally expressed in uniform inducing conditions. (A) Schematics showing SPI-2 activation inside host cells, SPI-2 genomic locus, and SPI-2 transcriptional reporter plasmid. Genes for which promoter-reporter constructs were created are shaded in green. Known transcription start sites are represented by arrows and a putative start site is shown with a dashed arrow. (B) Imaging of SPI-2 reporter strains shows bimodal expression in MgM-MES and unimodal expression in M9/Glc/CAA. PssaG (i) and PssaB (ii) reporter strains were grown in MgM-MES (inducing) or M9/Glc/CAA (non-inducing) for four hours, fixed, and imaged. Representative images for mRuby2, sfGFP and the merged channels are shown, with mRuby2 shown as magenta and sfGFP shown as green in the merged image. sfGFP intensity is scaled by reporter, mRuby2 intensity is scaled by media condition. Scale bars =10 µm. Density plots of single-cell intensity measurements of microscopy data from three independent experiments are shown. KDE contour plots of single-cell sfGFP intensity vs. mRuby2 intensity are shown. For all quantification n=450 cells per experiment, n=1350 per plot. (C) smFISH confirms bimodal expression of an endogenous SPI-2 gene. Representative images of WT STm grown in MgM-MES or M9/Glc/CAA for 4 hours and probed for *ssaL* transcripts. The percentage of cells with spots detected and the CDF of spots detected within cells (n=1200 cells, 300 per experiment), and the spot intensities for all intracellular spots are shown. (D) Samples evaluated via microscopy in (B) were analyzed via flow cytometry. Histograms from one representative experiment are shown. 50,000 cells per sample are displayed. Percentage GFP(+) cells as gated by the GFP(-) control is shown for three independent experiments.

While the environmental stimuli, gene regulatory network, and functional significance of SPI-2 have been well defined, SPI-2 activation in response to the intracellular environment poses several critical regulatory challenges for *Salmonella* that remain unexplored. First, phagosome environments can be variable in pH and nutrient content and change as the phagosome matures over the course of infection (Rathman et al., 1996; Drecktrah et al., 2006; Martin-Orozco et al., 2006; Dragotakes et al., 2020), forcing SPI-2 to reliably respond to a broad range of signals. Second, to survive host cell neutralization, a timely SPI-2 response is required despite some phagosomes containing suboptimal stimuli, creating frequent opportunities for failed responses. Third, the signals known to activate SPI-2 are not exclusive to the intracellular environment, particularly across the landscape of the entire gastrointestinal tract. While SPI-2 is essential for intracellular replication in tissue culture models and systemic infection in animal models (Fields et al., 1986; Cirillo et al., 1998; Han et al., 2024), loss of negative regulators of SPI-2 have been shown to impede pathogenicity, suggesting that aberrant responses to stimuli have a fitness cost during infection (J. Choi et al., 2010; Coombes et al., 2005). To what extent individual STm can discriminate between the intracellular environment and similar extracellular conditions while maintaining accurate responses to diverse phagosomes is unclear. Additionally, several previous studies report heterogeneity in SPI-2 expression both in host cells and *in vitro* (Cirillo et al., 1998; Hautefort et al., 2003; Blair et al., 2013; Stapels et al., 2018; Reuter et al., 2021). However, how such heterogeneity in SPI-2 expression originates or how subpopulations change in response to environmental stimuli remains unknown.

Here, we evaluated whether STm may have evolved a SPI-2 regulatory network that can tune transcriptional responses to intracellular stimuli at the single-cell level. We demonstrate that in *in vitro* inducing conditions, the bacterial population bifurcates into highly expressing and non-expressing subpopulations. Over time, individual cells switch into the high expressing state, but the bimodal distribution of gene expression is maintained across the population. We show that the rate of this transition can be altered by the strength and integration of intracellular-like signals, including decreases in pH and magnesium. We find that the regulation of the fraction of SPI-2 high expressing cells is gated by the concentration of the SPI-2 master regulator SsrB and established through SsrA-dependent signaling. Further, we show that this concentration based mechanism for temporally-delayed heterogeneity may allow for growth rate dependent filtering of intracellular-like signals. Lastly, we use live-cell imaging to robustly characterize the long term dynamics of SPI-2 activation *in vitro*, finding a committed switch-like transition that is stochastic in nature and correlated between closely related cells. We propose that the SPI-2 virulence system has been evolutionarily optimized to deal with environmental uncertainty by probabilistically tuning the likelihood of transcriptional responses to intracellular signals across time.

## RESULTS

### SPI-2 gene expression is bimodal under uniform inducing conditions

Because conditions in host cell phagosomes are highly dynamic and heterogeneous, we reasoned that an *in vitro* culture system using fluorescent transcriptional reporters would be ideally suited to investigate how individual STm regulate SPI-2 gene expression in response to intracellular-like signals. Our approach provides precise environmental control and enables accurate measurements of SPI-2 gene expression in individual bacteria, allowing us to directly link environmental inputs to gene expression outputs. To measure the transcriptional activity of the SPI-2 system in individual bacteria we designed promoter-based transcriptional reporters (Figure 1A). We focused on two major SPI-2 operons, selecting extensively characterized promoters upstream of *ssaG* (PssaG) and *ssaB* (PssaB) that have been used in similar reporter-based assays (Hautefort et al., 2003; Drecktrah et al., 2006; Tomljenovic-Berube et al., 2010; Osborne & Coombes, 2011; Luk et al., 2021). To preserve endogenous SPI-2 gene regulation, we selected a low-copy (pSC101) plasmid in lieu of disrupting the SPI-2 genomic locus. Our use of a low-copy plasmid also minimizes plasmid copy number variation and metabolic burden on the bacteria (Figure S1.1). To minimize the potential impact of fluorescent protein accumulation, we fused the *ssaG* promoter to a destabilized sfGFP (sfGFP-LVA) (Andersen et al., 1998), while the *ssaB* promoter was fused to stable sfGFP as the promoter was too weak to produce signal when fused to sfGFP(LVA) (Figure S1.2A). We included a constitutively expressed mRuby2 in our reporter plasmids to enable cell identification in microscopy and flow cytometry.

Using our SPI-2 reporter strains we examined SPI-2 reporter expression in response to pH 5.0 MgM-MES media, a limited medium that has been commonly used to mimic the phagosome environment and induce SPI-2 (Beuzón et al., 1999; Yu et al., 2004). We diluted each reporter strain (PssaG and PssaB) into SPI-2 inducing media (MgM-MES) or non-inducing media (M9/Glc/CAA) and quantified SPI-2 reporter expression in single cells after four hours using fluorescence microscopy. As expected, non-inducing conditions showed no reporter expression (Figure 1B). In contrast, both reporters showed bimodal expression patterns in SPI-2 inducing conditions, with distinct subpopulations of expressing and non-expressing cells (Figure 1B). This bimodality is independent of plasmid-copy number effects, as shown by unimodal mRuby2 expression that did not correlate with sfGFP signal in individual cells (Figure 1B). This is consistent with prior observations of SPI-2 heterogeneity (Cirillo et al., 1998; Hautefort et al., 2003; Blair et al., 2013; Stapels et al., 2018; Reuter et al., 2021) and demonstrates that there is intrinsic variation in SPI-2 activation given a uniform inducing environment.

Having established that SPI-2 reporter expression is bimodal, we next wanted to confirm that this bimodality reflected endogenous SPI-2 gene expression. We performed single-molecule fluorescence *in situ* hybridization (smFISH) for *ssaL*, the largest gene in the *ssaG* operon. We validated that the smFISH protocol accurately detects endogenous *ssaL* using Δ*ssaL* and Δ*ssrB* strains (Figure S1.3A) (Porwollik et al., 2014). Wildtype (WT) cells showed bimodality in transcript abundance in inducing conditions and little to no expression in non-inducing conditions (Figure 1C). Spot intensity in the induced samples was also greater than in non-induced samples, possibly reflecting nascent transcription (Figure 1C). Using the PssaG reporter strain we examined the relationship between endogenous SPI-2 transcription, by smFISH for *ssaL*, and GFP fluorescence. The percentage of cells expressing endogenous transcript and the reporter are similar and increased spot count trends with increased reporter activation (Figure S1.3B). However, overlap of the reporter and smFISH signal had discrepancies consistent with the expectations that 1) sfGFP(LVA) maturation follows *ssaL* transcription and 2) *ssaL* mRNA is less stable than the sfGFP(LVA) protein.

To enable higher throughput analysis of SPI-2 responses, we validated our microscopy results using flow cytometry. Both reporters showed consistent bimodal activation patterns across methods (Figures 1B, D), while mRuby2 expression remained unimodal (Figure S1.4), confirming that flow cytometry could be used to reliably quantify SPI-2 heterogeneity. We generated fluorescent protein variations of the reporter and evaluated induction by flow cytometry to confirm that heterogeneity was not due to sfGFP pH sensitivity (Figure S1.2 B-C). Lastly, we validated that the reporter was reflecting canonical activation through the known SPI-2 pathway using a knockout of the *ssrB* gene (Δ*ssrB*) (Figure S1.5A, C). Because PssaG was brighter than PssaB and could be used in combination with a destabilized sfGFP to limit accumulation based effects, we elected to use it as a representative SPI-2 reporter in further experiments. Overall, our results establish that STm SPI-2 expression is intrinsically heterogeneous in uniform environments and demonstrate that our reporter system is a suitable tool to interrogate the origins of this heterogeneity through both imaging and flow cytometry.

### SPI-2 activation increases over time through continuous single-cell transitions

Previously published examples of virulence heterogeneity show bistable expression, with discrete subpopulations of virulence expressing and non-expressing cells (Hautefort et al., 2003; Nielsen et al., 2010; Saini et al., 2010; Nuss et al., 2016; Ronin et al., 2017). We thus hypothesized that SPI-2 expression could either similarly be limited to a defined fraction of cells capable of responding to a given environment or that all cells have a capacity to respond to environmental signals given enough time. To test this, we cultured the PssaG reporter strain in inducing media and analyzed SPI-2 expression by flow cytometry at 2, 4 and 6 hours. Rather than maintaining a fixed ratio of expressing to non-expressing cells, we observed a progressive increase in the fraction of SPI-2 expressing cells over time (Figure 2A). This switch occurred over several cell generations based on growth rates in MgM-MES (Figure S1.1), indicating a delayed response to SPI-2 inducing conditions. To evaluate if this pattern was reflective of progressive transcriptional activation of the SPI-2 system rather than continuous accumulation of the reporter, we examined the fluorescence intensity of the two subpopulations over time. There is a SPI-2(-) population across all timepoints that maintains a consistent low intensity, while the positive population showed increasing intensity over time (Figure 2A). This pattern indicates the increase in SPI-2(+) cells is not likely to be due to accumulation and further suggests that cells that switch into a transcriptionally active state continue expressing the reporter throughout the experiment resulting in the increase in GFP intensity.

**Figure 2.**
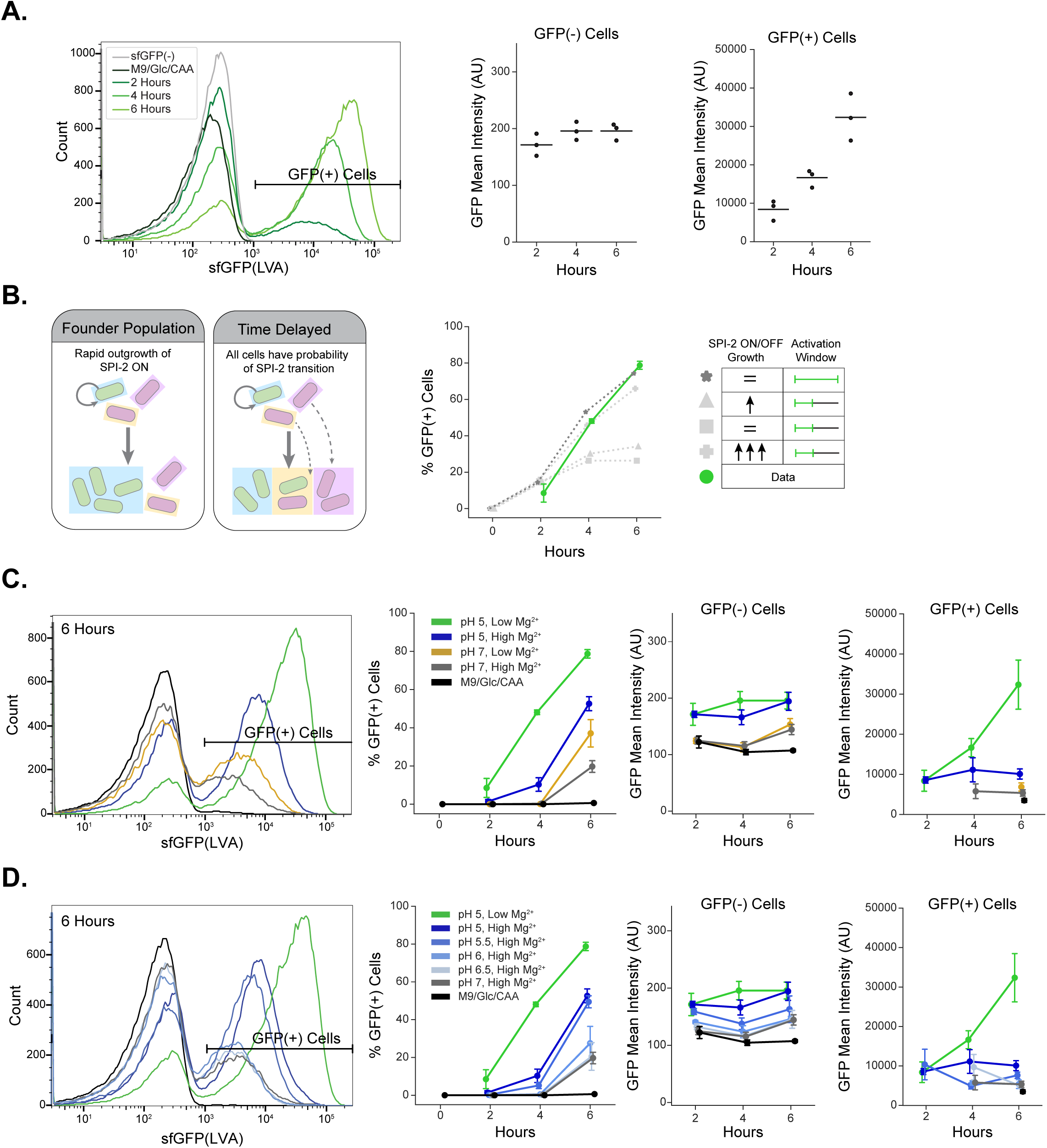
Single-cell SPI-2 activation is temporally-tunable to the strength and complexity of inducing signals. (A) Histograms of SPI-2 reporter strain in MgM-MES from a representative flow cytometry experiment show bimodality is maintained over time and the fraction of GFP(+) cells increases over time. Representative histogram of populations at 2, 4, 6 hours. n=50,000 cells per sample. Mean intensities for the GFP(+) and GFP(-) populations are displayed, n=3 experiments. (B) Comparison of two possible hypotheses for the generation of the bimodal distribution. In a ‘Founder Population’ model, a discrete fraction of cells turns on the reporter (green) at early time points and increase in frequency due to a faster growth rate than the negative population (pink). In a ‘Time Delayed’ model, all cells have a certain probability of activating the reporter for the duration of the experiment, and new SPI-2(+) cells continue to arise throughout the timecourse. Results of stochastic simulations in which the division rate of the two subpopulations (SPI-2(+) and SPI-2(-)) or the activation time window in which SPI-2 can be activated were altered. Only two simulation conditions match the experimental data (green): a time delayed condition with equivalent growth rates of the two subpopulations (*) or a founder population condition with the growth rate of the positive population set to an unrealistically faster rate than the negative subpopulation (+). Full parameters used are in Figure S2.1. (C, D) Comparison of SPI-2 induction rates in media with varied inducing signals (Mg^2+^, pH). Representative histograms from flow cytometry analysis at the 6-hour time point show that the rate of cells transitioning from SPI-2(-) to SPI-2(+) increases with multiple signals (C) and as the strength of an individual signal, pH, increases (D). n=50,000 cells per sample, n=3 experiments, t-tests in Table S1. The percentage of GFP(+) cells per time point and the mean intensities of the GFP(+) and GFP(-) populations are shown.

The gradual increase of SPI-2 positive cells could be explained in two possible ways: 1) a ‘Founder’ population model in which the increase of the SPI-2(+) population is due to the outgrowth of a population of initial responders or 2) a ‘Time-Delayed’ activation model in which cells continuously transition into a SPI-2 active state throughout the experiment (Figure 2B). To distinguish between these models, we used simple simulations of fixed probability transitions with different subpopulation growth rates and SPI-2 activation windows. We show that our experimental data is most likely consistent with cells entering a SPI-2 transcriptionally active state *de novo* rather than a discrete population of responders giving rise to the SPI-2(+) cells via replication (Figure 2B). A ‘Founder’ model would require SPI-2(+) cells to grow approximately five times faster than SPI-2(-) cells to match our experimental data (Figure 2B and Figure S2.1), an unrealistic difference given our growth data (Figure S1.1). In contrast, the ‘Time-Delayed’ model reproduced our experimental data using realistic growth parameters. These results demonstrate that SPI-2 activation is a continuous process, with individual cells progressively transitioning into a SPI-2 expressing state over time, rather than a bistable system with defined responders and non-responders.

### Intracellular-like signals tune the rate at which SPI-2 is activated

As phagosomes are known to vary in both magnesium concentration and pH (Rathman et al., 1996; Drecktrah et al., 2006; Martin-Orozco et al., 2006; Dragotakes et al., 2020), we hypothesized that SPI-2 activation might be tuned to the strength of multiple intracellular-like signals, enabling a flexible response to heterogeneous environmental conditions. To test this, we compared PssaG reporter activation across different pH levels (5.0 - 7.0) and magnesium concentrations (8 µM vs 2 mM) over 6 hours by flow cytometry. We find cells cultured in the strongest inducing condition (pH 5.0, 8 µM Mg^2+^) have the most rapid transition to a SPI-2 expressing state, while weaker inducing signals (i.e. pH 5 alone) slowed the rate at which cells transitioned into the SPI-2 expressing state (Figure 2C). Strikingly, all inducing conditions maintained bimodal expression patterns (Figure 2C and S2.2). pH appears to be the dominant signal in our experimental conditions, as acidic conditions alone produced rapid SPI-2 reporter activation while low magnesium alone did not significantly induce reporter activation until 6 hours of growth (Figure 2C). Therefore, to test whether the rate of transition to the SPI-2 expressing state could be tuned by the strength of a single input signal, we compared SPI-2 activation rates in response to incremental changes in pH (Figure 2D). SPI-2 activation only occurred prior to the 6 hour time point when the pH was <= 6, with no significant differences between cells grown in pH 5 or pH 5.5 media, or between pH 6 - 7, despite expected decreases in growth with decreases in pH (Figures 2D and S2.3). This sharp pH threshold is consistent with recent biochemical studies showing that the SsrB protein undergoes a conformational change at a pH <5.8 that increases DNA binding affinity (Liew et al., 2019; Shetty & Kenney, 2023). All together, these results suggest that the rate of SPI-2 transitions in single cells increases with stronger intracellular-like signals. This rate-based tuning mechanism allows for a fractional response within a given time window in the event of an ambiguous environment containing relevant intracellular signals.

### The master regulator SsrB is critical for bimodality in SPI-2 expression

We next sought to understand how the regulatory network of SPI-2 generates this tunable, time-delayed heterogeneity. The SPI-2 network has been well defined and a simplified schematic is shown in Figure 3A. We tested knockout strains of major SPI-2 regulators (Porwollik et al., 2014) using our PssaG reporter system, including *phoP*, *ptsN*, *fur*, and *ssrB*. SPI-2 activation was assessed using flow cytometry. We found that deletion of negative regulators, *ptsN* and *fur,* increased the rate at which individual cells transition into the SPI-2 expression state without eliminating the bimodality in expression or allowing aberrant expression in non-inducing conditions (Figure 3B and S3.1). However, both of these mutations also reduce growth in MgM-MES (Figure S3.2), limiting our ability to distinguish between regulatory and accumulation-based effects. Overall, these data show that negative regulators alter the rate of SPI-2 activation, but otherwise maintain the bimodal distribution and the progressive single-cell switch into the SPI-2(+) state in inducing conditions

**Figure 3.**
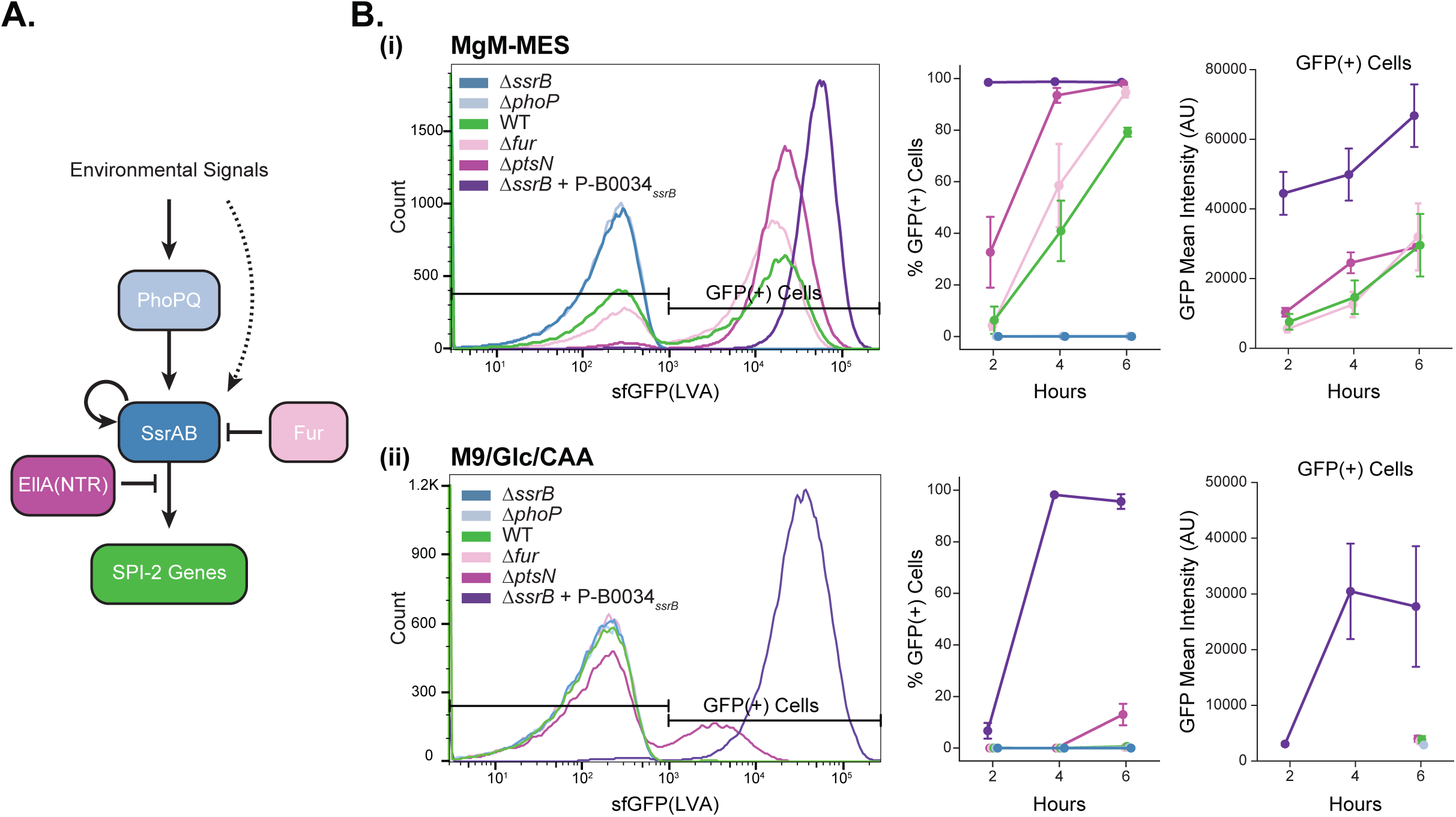
Tunability of SPI-2 activation depends on the expression of *ssrB* while the timing of activation is tuned via known negative regulators. (A) Simplified schematic of known SPI-2 regulators including upstream two-component systems (PhoPQ, SsrAB) and negative regulators (EllA(NTR), Fur). (B) To evaluate which known SPI-2 regulators are necessary for bimodality in SPI-2 expression, the PssaG reporter was transformed into knockout strains of these known regulators or co-transformed with a constitutive SsrB expressing plasmid (P-B0034*_ssrB_*). Strains were then grown in MgM-MES (i) or M9/Glc/CAA (ii) and analyzed via flow cytometry for reporter expression at indicated timepoints. A representative histogram at four hours (n=50,000 cells/sample), the percentage of GFP(+) cells, and the mean intensities of the GFP(+) population are shown for each medium across timepoints (n=3 experiments, t-tests in Table S1).

As expected, our results further show that loss of *ssrB* or upstream two-component systems (*phoP*) completely abolishes SPI-2 reporter expression (Figure 3B). Given the central role of *ssrB*, we hypothesized that SsrB abundance may control SPI-2 heterogeneity. To test this hypothesis, we complemented the *ssrB* knockout strain with a plasmid constitutively expressing *ssrB* at relatively low levels (P-B0034*_ssrB_*, see Methods) along with the reporter plasmid (Figure 3B). Attempts at expressing SsrB at levels higher than this construct were toxic. SsrB expression from the P-B0034*_ssrB_* plasmid changed expression from bimodal to near unimodal for both the PssaG and PssaB reporters in inducing and non-inducing conditions (Figure S1.5B), with *ssaL* transcripts detectable by smFISH in almost all cells (Figure S1.3A). The exception occurred at 2 hours in M9/Glc/CAA with <10% of cells expressing the reporter (Figure 3B), which we hypothesize is due to rapid growth diluting plasmid-expressed SsrB below a critical concentration threshold needed to activate SPI-2 genes. Combined with the observation that there is no expression of the reporter in the absence of SsrB or upstream activators, these results indicate that heterogeneity in endogenous SPI-2 expression is dependent on a low expression level of SsrB and defined in part by SsrB levels.

### Heterogeneity in SPI-2 expression depends on both SsrB concentration and SsrA signaling

Because overexpression of *ssrB* produces near unimodal expression of SPI-2 (Figure 3B), we hypothesized three possible SsrB-dependent mechanisms by which heterogeneity in SPI-2 expression could be established. The first is a production-dependent delay mechanism, in which stochastic variation in the timing of *ssrB* gene expression results in heterogeneity in the timing of SPI-2 expression. This heterogeneity could arise from variation in upstream two-component systems sensing the environmental shift or variation in basal SsrB expression levels. The second is an autoregulation-dependent mechanism. Since positive feedback is a common regulatory feature in bimodal systems (Saini et al., 2010; Smits et al., 2006) and SsrB has a putative binding site in its own promoter (Tomljenovic-Berube et al., 2010), this could lead to bifurcation of the population at low SsrB concentrations (Figure 4A). Third is a signaling-dependent mechanism. SsrB concentration is normally distributed across the population, but SsrA signaling is bimodal, resulting in the formation of two distinct subpopulations (Figure 4B).

**Figure 4.**
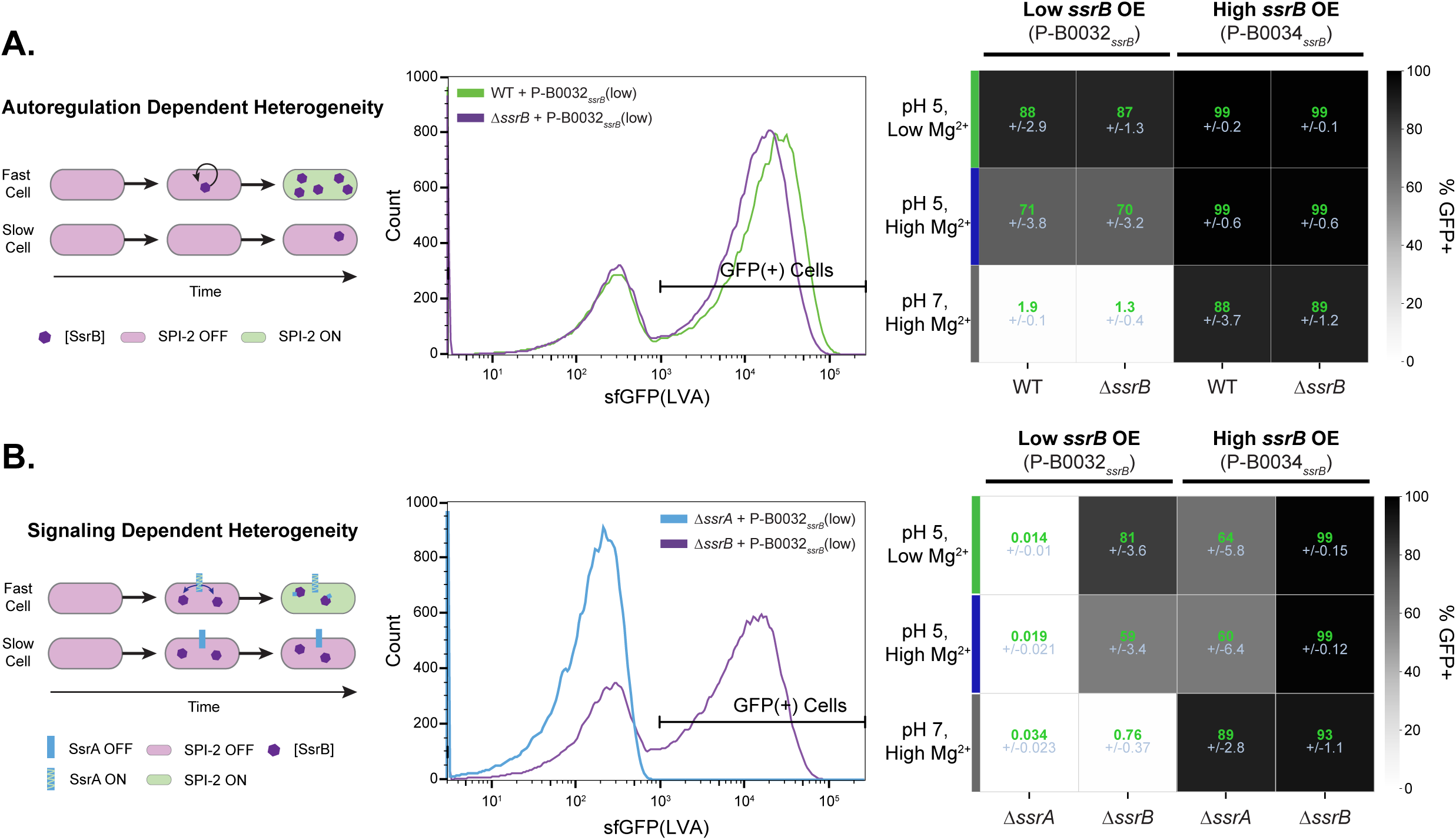
Heterogeneity in SPI-2 expression is dependent on both the concentration of SsrB and signaling through SsrA. We tested the mechanism of SsrB-dependent heterogeneity using low, constitutive expression of SsrB (P-B0032*_ssrB_*). (A) In an autoregulation-dependent mechanism, stochastic noise in *ssrB* expression is amplified via positive feedback, promoting the bimodality in SPI-2 expression (left). A representative histogram of PssaG reporter signal in the WT and Δ*ssrB* strains containing P-B0032*_ssrB_* grown in pH 5, 2 mM Mg MgM-MES. Both strains are similarly bimodal (middle). A heatmap of % GFP(+) cells across conditions (Low Mg_2+_ = 8 µM, High Mg_2+_ = 2 mM) and ssrB expression constructs based on flow cytometry after 4 hours of growth shows there are no significant differences between a WT and Δ*ssrB* strain across conditions (right, t-tests in Table S1). (B) In a signaling-dependent model, variation in SsrA signaling promotes the bimodality in SPI-2 expression (left). A representative histogram of PssaG reporter signal in the Δ*ssrA* and Δ*ssrB* strains containing P-B0032*_ssrB_* grown in pH 5, 2 mM Mg MgM-MES. The Δ*ssrA* strain has no expression of downstream genes (middle). A heatmap of % GFP(+) cells across conditions and ssrB expression constructs based on flow cytometry after 4 hours of growth (right) shows significant differences between the Δ*ssrA* and Δ*ssrB* strains containing P-B0032*_ssrB_* (low *ssrB* OE).

Expressing uniform low, constitutive levels of SsrB below the threshold needed for unimodal SPI-2 activation would allow us to systematically test each hypothesis by evaluating reporter expression patterns in the absence of endogenous SsrB variation. To accomplish this, we generated a second *ssrB* expression plasmid that produces lower SsrB levels than our original plasmid. Since the promoter and copy number of the original plasmid (P-B0034*_ssrB_*) were already weak (Methods), we used a weaker synthetic ribosome binding site, BBa_B0032 (Smith et al., 2023) to further reduce expression. We validated that changing the RBS decreases protein production (Figure S4.1). While we initially aimed to directly correlate SsrB levels with reporter output, we were unable to tag SsrB without altering reporter activation (Figure S4.2). We thus elected to evaluate reporter output to untagged SsrB expressed at the lower concentration. Using flow cytometry, in non-inducing conditions (pH 7, 2 mM Mg^2+^), reduced SsrB levels did not activate the PssaG reporter in the Δ*ssrB* strain in contrast with the original P-B0034*_ssrB_* plasmid, which promoted uniform activation in these conditions (Figure 4A). However, in inducing conditions reporter bimodality was still observed in strains with the P-B0032*_ssrB_* plasmid similar to endogenous activation, suggesting that heterogeneity in SsrB production or levels (hypothesis #1) alone is not responsible for heterogeneity in SPI-2 (Figure 4A, middle). We further compared reporter output in the WT and Δ*ssrB* strains containing P-B0032*_ssrB_* across a range of conditions and timepoints to evaluate whether SsrB autoregulation (hypothesis #2, Figure 4A and S4.3A) is required for bifurcation of the two subpopulations. We found no significant differences, suggesting that autoregulation is not responsible for heterogeneity in SPI-2 expression. The high overexpression construct promoted high unimodal reporter signal and was likewise unchanged in the presence or absence of the endogenous *ssrB* gene (Figure 4A).

To test whether an SsrA signaling dependent mechanism is responsible for bimodal expression (hypothesis #3, Figure 4B) we compared reporter output between the Δ*ssrA* and Δ*ssrB* strains containing the low *ssrB* overexpression plasmid (P-B0032*_ssrB_*) . At low SsrB levels in the Δ*ssrA* strain, SPI-2 reporter signal was completely abolished (Figure 4B). This indicates that given equal concentration of the response regulator SsrB, there is bimodal variation in either SsrA signaling capacity or concentration in inducing conditions, leading to variation in SsrB binding to downstream genes and bimodal SPI-2 expression. We further evaluated reporter output in these strains with the higher *ssrB* overexpression plasmid and found that in an Δ*ssrA* background, high SsrB concentration alone generated unimodal reporter output in non-inducing media, though there were unexpected, likely dilution-based effects in inducing media (Figure 4B and S4.3B). Together, these results support a model in which heterogeneity in SPI-2 expression arises primarily at the level of SsrA signaling when SsrB is present at endogenous levels. However, the fraction of cells expressing SPI-2 can still be controlled by increased SsrB concentration alone in the absence of SsrA signalling.

### Environmental modulation of growth rate sensitizes cells to SPI-2 inducing signals

Because our results demonstrate that SPI-2 activation is sensitive to SsrB concentration, we next explored how global dilution of molecules via cell division might impact the system (Klumpp et al., 2009; Kueh et al., 2013; Klumpp & Hwa, 2014; Narula et al., 2016; Zatulovskiy et al., 2020). Recent work has demonstrated through modeling and experiments that modulation of growth rate alone can sensitize or desensitize bacterial responses via accumulation or dilution of regulatory proteins (Julou et al., 2025). We therefore hypothesized that the SPI-2 regulatory network might be sensitive to environmentally induced growth perturbations because they alter the rate of accumulation of SsrA and SsrB (Figure 5A). We first tested this hypothesis by modulating growth rate through carbon source variation in pH 5 MgM-MES minimal media, the strongest inducing condition from prior experiments (Figure 2C). Glucose increased the growth rate of the population relative to the standard glycerol condition, while pyruvate decreased the growth rate (Figure S5.1). We found that the percentage of responding cells increased with decreasing growth rate, despite identical SPI-2 inducing signals, with the most significant differences observed at the 2-hour time point (Figure 5B). We next investigated whether changes in growth rate could sensitize a population to a weaker SPI-2 inducing signal. To test this, we used a pH 7, low Mg^2+^ medium with casamino acids (CAA), as in this condition cells do not respond until 6 hours and growth can be substantially reduced by removing CAAs (Figures 5B and S5.1). While in the CAA-replete condition there were no SPI-2 responders at four hours of growth, the absence of CAA resulted in a significant increase in responders under these conditions (Figure 5B). Notably, this effect did not occur in a CAA-limited condition that lacked the low magnesium stimulus (Figure 5B). These results further support the hypothesis that the rate of the SPI-2 response can be altered by growth rate and demonstrate that this effect is not specific to carbon metabolism. In both experimental conditions, we observed that the fluorescence intensity of the ‘OFF’ population was maintained under slower growth conditions, suggesting that the increased ‘ON’ population reflected a transition into a transcriptionally active state rather than reporter accumulation (Figure S5.2). To conclusively validate that the increase in responders reflected SPI-2 activation and not GFP accumulation, we performed smFISH on *ssaL* transcripts under representative slow growth conditions. We found that endogenous transcripts were increased in slow growth conditions relative to the identical SPI-2 stimuli in faster growth conditions (Figure 5C and S5.3A). Together, our results suggest that growth rate may modulate SPI-2 response timing to equivalent intracellular stimuli, revealing additional complexity in the environmental regulation of the system.

**Figure 5.**
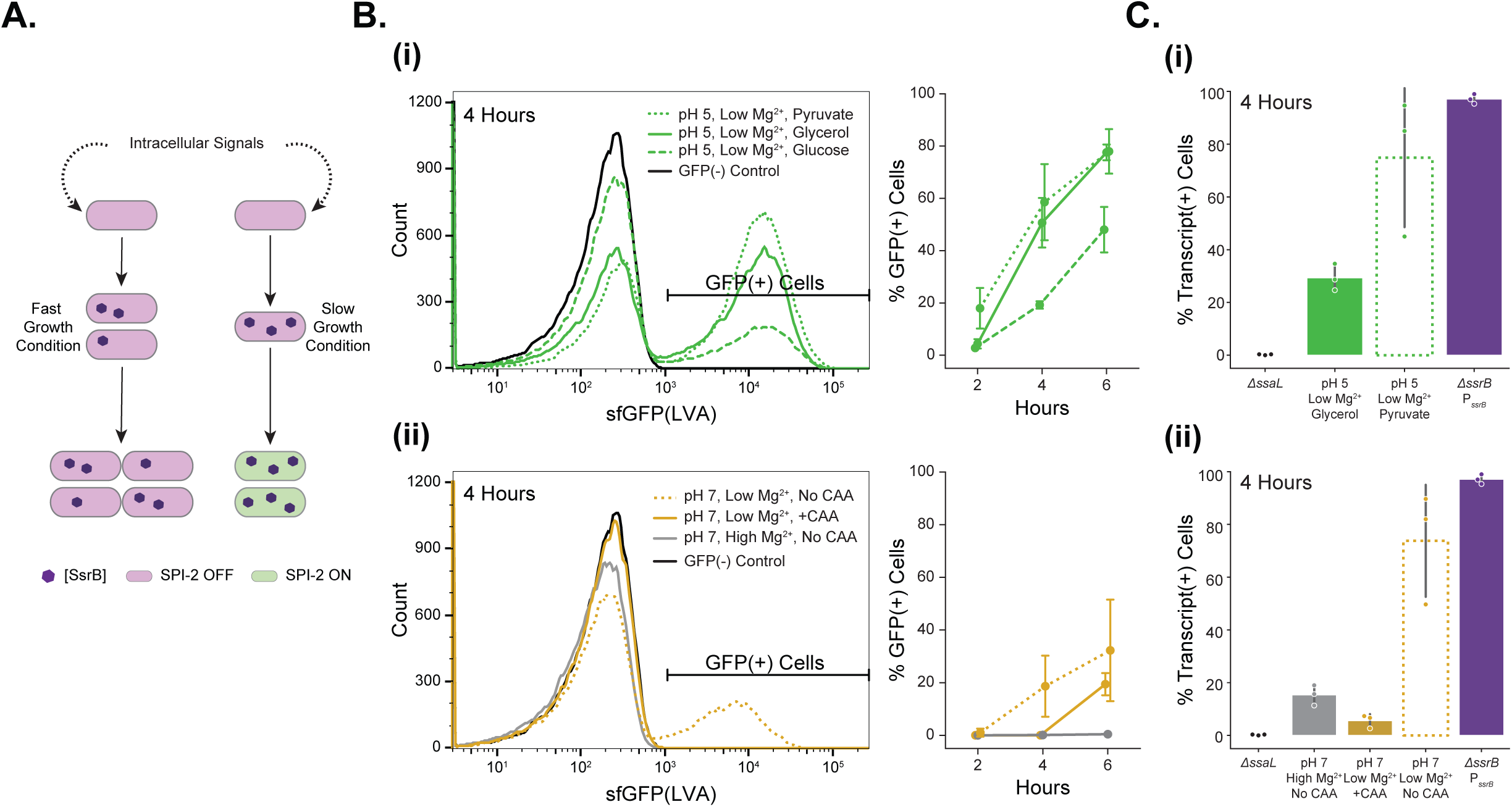
Changing nutrient content to alter growth rates changes the rate at which cells activate SPI-2. **(A)** Schematic illustrating the hypothesis that slowing growth of cells may sensitize them to equivalent intracellular signals through SsrB or other regulatory factor accumulation. (B) To test this hypothesis, cells containing the SPI-2 reporter were grown in strong inducing conditions (pH 5, 8 µM MgCl_2_ MgM-MES) with varied carbon sources to alter growth rate and analyzed via flow cytometry (i). Cells containing the reporter were also grown in weak inducing conditions (pH 7, 8 µM MgCl_2_ MgM-MES) with and without casamino acids (CAA) and analyzed via flow cytometry (ii). For both strong and weak inducing conditions, a representative histogram of reporter intensity after 4 hours is shown (n=50,000 cells per sample) alongside the percentage of GFP(+) cells across time in the conditions (n=4 experiments, error bars = standard deviation, t-tests in Table S1). (C) To evaluate whether the results with the reporter are reflected at endogenous SPI-2 genes, smFISH was performed using *ssaL* transcript probes in select conditions with carbon variations in strong inducing conditions (i) or CAA variations in weak inducing conditions (ii) at 4 hours. The percentage of cells with transcripts detected are shown. Slower growth conditions increase the fraction of cells responding at both the reporter and endogenous transcript level.

While our results support a link between growth rate and SPI-2 responses, several caveats limit these findings including that 1) these nutrient modifications are harsh and likely affect cell metabolism and stress responses beyond altering growth rate; 2) we observed increased variability in SPI-2 reporter induction under extremely slow-growing conditions (Figure 5B); and 3) smFISH results showed higher induction at both the endogenous transcript and reporter signal levels in the no CAA, neutral pH, low magnesium condition relative to pH 5 MgM-MES (Figure S5.3). Despite these confounding results, the data for both the reporter and endogenous transcript levels support our hypothesis that given equivalent SPI-2 inducing signals, environmental changes that reduce growth rate increase the rate at which the population transitions to a SPI-2 active state. While these results are perhaps not surprising given the system’s dependence on low numbers of regulatory proteins, this presents an additional mechanism by which single cells could distinguish the phagosome environment from extracellular environments.

### SPI-2 activation is switch-like and committed at the single-cell level

Our results indicate that STm tunes the rate of single-cell SPI-2 transitions to its environment, however, these conclusions are limited by our use of fixed time point observations and a closed culture system where the environment changes as cell density increases. Cellular responses to environmental transitions can also be tuned at the single-cell level across time and capturing the trajectory of the single-cell response is critical to link heterogeneity with functional outcomes (Patange et al., 2018; Sampaio et al., 2022; Spratt & Lane, 2022). To characterize activation of the SPI-2 system without such limitations and resolve the dynamics and stability of single-cell responses, we used the Dual Input Mother Machine (DIMM) (Kaiser et al., 2018) to monitor growth and SPI-2 reporter expression in real time following the transition into a SPI-2 inducing environment (Figure 6A). Cells were grown and imaged in M9/Glc/CAA (non-inducing media) for at least 5 hours, then transitioned to standard MgM-MES (inducing media) and imaged every 5 minutes for 18 hours. As expected, no reporter activation was observed during growth in M9/Glc/CAA (Figure S6.1). Following the transition into MgM-MES, we quantified SPI-2 reporter signal and found characteristic bimodality. The SPI-2(+) population gradually increases over time after a delay of several hours in which no cells respond, consistent with our observations in the closed culture system (Figures 2A, 6B-C). The mRuby2 intensity of the population remains fairly constant across time and shows no correlation with reporter activation (Figure S6.2)

**Figure 6.**
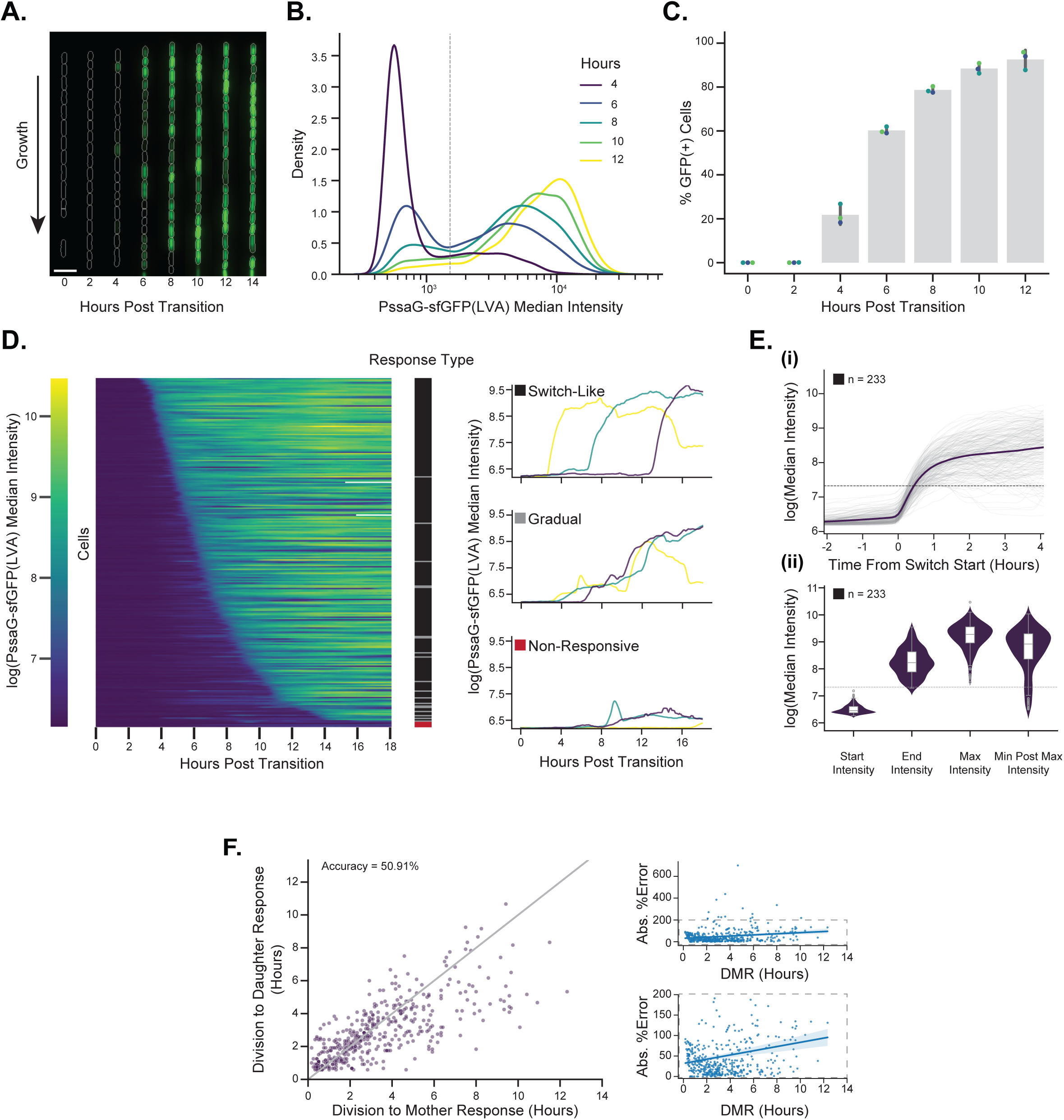
Single-cell trajectories reveal dynamic and heritable features of SPI-2 activation. (A) A representative kymograph of a single lineage of cells containing the PssaG-sfGFP(LVA) reporter after the transition from M9/Glc/CAA to MgM-MES media in the DIMM is shown. Each frame is equal to 2 hours, scale bar = 5 µm. (B) Density plot of all cells in the DIMM across time shows bimodality in reporter expression. n>4900 cells for each timepoint. Cells are pooled from three independent experiments. The dashed line represents the threshold used for calling cells as GFP(+) (Methods). Hours are post transition into MgM-MES media. (C) Percentage of GFP(+) cells increases across time. Each point represents data from an independent experimental replicate. (D) SPI-2 responses are predominantly switch-like, with cell-to-cell variation in time to respond. Mother cells (n=258, pooled from three experiments) were sorted by the time the cell crosses the GFP(+) intensity threshold. Time series data of mother cell PssaG-sfGFP(LVA) reporter intensity is plotted as a heatmap with each row representing a single mother cell and the columns representing time points. Mother cells were classified into one of three response types (switch-like (black), gradual (gray), or non-responsive (red)) based on features of the response. The response type of each mother cell is shown in the color bar to the right of the heatmap. Representative traces of reporter intensity within each response type are displayed. (E) Trajectories and features of the response in switch-like mother cells (n=233). (i) Switch-like mother cell PssaG-sfGFP(LVA) reporter intensities normalized to the time of start increase as defined by the slope of the intensity over time (Fig. S6.4) (top). (ii) Intensity features of mother cell switch responses across the experiment (bottom). (F) Correlation of mother and daughter cell reporter activation times relative to the time of division. Gray line represents y=x. Accuracy is equal to 100 - MAPE value relative to the y=x correlation. Absolute percent error of each parent cell start time relative to the y=x line across time (bottom is adjusted y-axis to visualize distribution of the population without outliers).

Having validated that SPI-2 bimodality and progressive activation was maintained in the DIMM, we next examined individual cell trajectories to the SPI-2 expressing state to determine transition rates and the stability of the response. We quantified the SPI-2 response in mother cells over the duration of the experiment and then sorted the data based on their time to cross the GFP(+) threshold (Methods). We observed a wide range in mother cell activation times (∼4-18 hours) (Figure 6D) with nearly all mother cells (>98%) eventually responding. Using the mother cell reporter time series data to calculate the SPI-2 ON and OFF rates, we found SPI-2 activation was committed, with a higher ‘ON’ rate (0.152/hr) than ‘OFF’ rate (0.027/hr) (Figure S6.3A-B). The probability of cells transitioning into a SPI-2 ‘ON’ state increases sharply after three hours of growth in inducing conditions, and remains stable from approximately four hours until 10 hours when the majority of the population is expressing SPI-2 (Figure S6.3C). These data confirm our ‘Time-Delayed’ model of SPI-2 activation (Figure 2). We further found mother cells predominantly (90.6%) activated SPI-2 in a rapid, switch-like manner (Figure 6D) as defined by the slope of the sfGFP(LVA) signal intensity across time (Figure S6.4, Methods). To compare the trajectories of the switch-like cells, we aligned them based on the time the reporter began to first increase and found their trajectories to the SPI-2 ON state were highly similar, demonstrating that the cells vary primarily in the time at which they initiated activation rather than the mode of activation (Figures 6D-E). Combined, our SPI-2 time series data reveal that SPI-2 activation is predominantly switch-like and committed at the single-cell level, with bimodality in the population arising from heterogeneity in the initiation of activation.

### SPI-2 response time is correlated between direct descendents

Having established that variation in the initial timing of activation is where SPI-2 heterogeneity arises, we next sought to understand whether this timing could be related to dynamic features we were able to measure during the DIMM experiments. We first evaluated whether timing of activation was strongly linked to pre-existing variation prior to the media transition by assessing SPI-2 activation on a per channel basis, as all cells within a channel originate from the same mother cell. We found variation within and across channels, with no obvious trend between the timing of initial responders in the channel and how quickly the entire channel activated SPI-2 (Figure S6.5A). We also confirmed that SPI-2 was not activated via a cell-cycle counter mechanism, as mother cells activated SPI-2 across a range of cell division events (Figure S6.5B). Another possible source of variation we considered was the rate of acidification of the STm cytoplasm, however recent measurements have shown this to be fairly uniform across populations (Singh et al., 2025). Given our observations that environmental manipulations of growth rate can alter the rate of SPI-2 activation (Figure 5), we investigated whether features of cell growth can predict the time of SPI-2 activation. We measured cycle duration, length at division, and the number of progeny produced by each lineage and observed considerable variability and irregularity in cell growth in SPI-2 inducing media (Figure S6.6). To determine if this variation contributed to the heterogeneity in SPI-2 activation timing, we trained a Random Forest Regressor (RFR) model on features of cell growth, mRuby2 expression, and experimental setup variables (Figure S6.7), an approach recently employed to identify determining features of the *E. coli* oxidative stress response (Choudhary et al., 2023). Our model was poorly predictive of SPI-2 initiation time. However, SHAP analysis of contributing features to the model identified that outliers in cell growth features corresponding to fast growth rates did predict longer delays in SPI-2 activation (Figure S6.7). These results are consistent with our hypothesis that division-based dilution of regulatory proteins can modulate activation of the system (Figure 5). The limited predictive power of this model indicates that SPI-2 activation is primarily driven by stochastic processes rather than cell growth features, but does not exclude growth as a contributing factor to response time.

Lineage analysis has found correlations between mother and daughter cell features (Ronin et al., 2017; Chatterjee & Acar, 2018; Lugagne et al., 2020; Vashistha et al., 2021), primarily due to the inheritance of long lived proteins across generations. We thus reasoned that the timing of SPI-2 activation could be linked between closely related cells. To test this, we examined SPI-2 response times for all mother-daughter pairs in which 1) the mother had not initiated SPI-2 activation before division, 2) both cells responded to the inducing media, and 3) the daughter cell responded 30 minutes or later following division. We normalized the start time of SPI-2 activation for each pair to the time of division. We find that mother and daughter SPI-2 response times are correlated (Figure 6F). However, substantial deviation from this relationship remained, with the prediction error increasing with longer time from division to initiation. These results confirm that SPI-2 activation is primarily stochastic, but suggest this stochasticity can be linked across cell divisions.

## DISCUSSION

Here, we used the *Salmonella* virulence system SPI-2 as a model for investigating environmental response tuning in bacteria. By using a reduced, *in vitro* experimental approach, we were able to evaluate input-output relationships in SPI-2 gene regulation in the absence of the inherent heterogeneity of host cell phagosomes (Rathman et al., 1996; Drecktrah et al., 2006; Martin-Orozco et al., 2006; Dragotakes et al., 2020). At the transcriptional level, we have shown that SPI-2 is bimodally expressed and individual cells progressively transition into a SPI-2 active state over multi-generational time periods. We found that the rate of this transition is modulated by intracellular-like signals and that this tunable bimodality is mechanistically dependent on low concentrations of the response regulator SsrB and heterogeneous signaling through the sensor kinase SsrA. Lastly, we demonstrated that the dynamics of SPI-2 activation are predominantly switch-like and committed, with heterogeneity arising from the initial timing of the response. We further show a correlation between the response times of mother-daughter pairs, indicating variation in response timing is partially heritable. Our results support a model in which the SPI-2 response of a clonal population of *Salmonella* is probabilistically tuned to the environment through varied single cell commitment rates, a strategy that may have evolved to resolve environmental uncertainty.

### Features of SPI-2 transcriptional tuning and their evolutionary implications

Our data reveal several unique features of SPI-2 tuning that may reflect evolutionary advantages to this kind of regulatory strategy. The first is that this system does not have a defined stimulus threshold above which a response is activated; even weak inducing environments are capable of activating a fractional SPI-2 response (Figure 2). The range of signals capable of inducing the system likely reflects the broad range of phagosome environments cells must respond to in order to replicate intracellularly. Environmental regulation of SPI-2 has been thoroughly investigated, with some conflicting data on what the primary phagosomal stimuli for the system may be (Alpuche Aranda et al., 1992; Rathman et al., 1996; Deiwick et al., 1999; Garmendia et al., 2003b; Löber et al., 2006). While our results favor pH being the predominant signal that activates SPI-2 *in vitro*, they also posit a mechanism for how multiple phagosomal signals can be integrated to tune responses through the rate of single-cell commitment. Future work could explore whether this rate-based tuning holds true for other signals that have been shown to modulate SPI-2 expression, such as calcium, iron, and phosphate levels (Deiwick et al., 1999; Garmendia et al., 2003; E. Choi et al., 2014).

Another defining feature of SPI-2 regulation this work demonstrates is that the majority of cells have the capacity to respond to phagosome-like signals (Figure 6). This points to SPI-2 phenotypic heterogeneity likely not being a division of labor based strategy, but rather a probabilistic strategy that will allow any phagocytosed bacteria to eventually respond to the intracellular environment given enough time. Previous observations of SPI-2 heterogeneity in cell culture infections (Cirillo et al., 1998; Hautefort et al., 2003; Reuter et al., 2021; Stapels et al., 2018) may thus be reflective of single-cell variations in the time to activate the system, or otherwise indicate there is a defined window of opportunity in host cells during which STm are capable of activating SPI-2. To what extent the timing of SPI-2 activation corresponds to single-cell infection outcomes such as phagosome maturation, bacterial replication, or host innate immune responses will require advances in intracellular tracking of individual bacteria. Because SPI-2 responses in the phagosome dictate bacterial survival (Cirillo et al., 1998; Fields et al., 1986; Han et al., 2024), the broad distribution in single-cell response sensitivities we’ve observed suggests a tradeoff for a guaranteed optimal response of a subset of individual bacteria in a range of phagosome environments at the expense of potential non-responders.

Conversely, the wide range in SPI-2 activation times may simply suggest an evolutionary prioritization of SPI-2 ‘OFF’ cells. Virulence expression is often described as energetically costly for cells (Davis, 2020). Long term maintenance of the SPI-2 ‘OFF’ population in inducing environments may have therefore evolved simply to offset the physiological resource requirements of expression of the T3SS and secreted effectors. However, virulence heterogeneity can have other benefits for bacterial populations beyond limiting energetic costs. The STm SPI-1 T3SS that is required for epithelial cell invasion, for example, is expressed bistably, and employs a unique division-of-labor strategy with both SPI-1(-) and SPI-1(+) cells required to promote gut colonization (Ackermann et al., 2008; Diard et al., 2013; Sánchez-Romero & Casadesús, 2018). Identifying the functional significance of single-cell variation in SPI-2 in regards to both bacterial physiology and population level infection success will help guide the understanding of how and why this rate-tunable system evolved.

Finally, the committed, switch-like dynamics of SPI-2 activation are significant findings for several reasons. Such commitment indicates the system lacks negative feedback mechanisms to halt expression under continuous exposure to inducing environments. It can be reasonably assumed then, that residence within a stable phagosome environment would sustain SPI-2 expression for as long as STm resides and replicates within the host cell. However, the complex dynamics of phagosome acidification and nutrient depletion and their dependence on STm activity (Drecktrah et al., 2006; Jennings et al., 2017) may alter this commitment in unexpected ways, making further evaluation of dynamic environmental inputs in similar *in vitro* experiments necessary. Switch-like activation in which heterogeneity across the population is primarily dependent on variation in activation time, suggests that there is limited tuning of SPI-2 transcription beyond the timing of an ‘OFF’ or ‘ON’ response. This binary tuning of the structural operons (PssaG and PssaB) is intuitive, perhaps, as the construction and embedding of a complex macromolecule like the T3SS is an all or nothing functional outcome, with only 1-2 SPI-2 injectisomes being observed on the surface of active cells (Chakravortty et al., 2005). While SPI-2 regulation appears deterministic at the level of the structural operons, multiplexed reporter experiments may reveal further complexities in distal components of the regulon, as has been observed in the *E. coli* flagella system (Kim et al., 2020).

### Mechanistic insights into SPI-2 regulation and origins of heterogeneity

Our results also contribute to the mechanistic understanding of how SPI-2 genes are induced, specifically in how SsrB protein levels correlate with downstream genes. In recent work where single-molecule imaging was performed on SsrB at endogenous levels, it was found that SsrB levels do not correlate with SPI-2 injectisome presence (Liew et al., 2019). Consistent with these results, our data show that synthetic expression of SsrB at low uniform levels generates a population heterogeneously expressing the PssaG reporter. Based on our findings in the Δ*ssrA* strain, we propose the disconnect between SsrB levels and downstream SPI-2 gene activation is due to heterogeneity in SsrA signaling in both of these cases. Future work could evaluate whether this heterogeneity is in SsrA levels or kinase activity, and whether feedback from SsrB on SsrA is necessary to generate this bifurcation.

Our finding that increasing the concentration of SsrB will generate a uniformly expressing SPI-2 population can likely be attributed to the fact that this synthetic expression is above endogenous levels, suggesting that the number of SsrB molecules is constrained to very low levels. This constraint may be advantageous as it allows for the generation of heterogeneity in SPI-2 expression in the population, and may further allow for more rapid deactivation of SPI-2 as cells transition out of an intracellular environment. These observations of high sensitivity to low concentrations of regulatory proteins are similar to model systems such as the *E. coli* lac operon, in which bistable activation is triggered by stochastic variation on the order of one molecule (P. J. Choi et al., 2008; Julou et al., 2020). We note that care must be taken in the interpretation of overexpression studies of SPI-2 for this reason, as single-cell heterogeneity, which appears to be a critical information processing feature of the system, is entirely masked above endogenous levels of SsrB.

We further show that this concentration sensitivity allows for modulation of SPI-2 response timing through environmental perturbations of growth rate (Figure 5). This phenomenon has been shown in synthetic systems and more recently the *E. coli* lac operon (Tan et al., 2009; Julou et al., 2020). Because initial divisions of STm within a macrophage in cell culture can take several hours, (Helaine et al., 2010) slowed growth rate within the phagosome may serve as an additional physiological cue in conjunction with pH and low magnesium. The integration of growth rate as an additional contextual signal could therefore more effectively filter cases of acid or nutrient deprivation stress occurring extracellularly. We propose that constraint of regulator proteins to low, deterministic concentrations may be a widely employed bacterial response strategy, as it allows for growth rate based modulation of responses, rapid transitions into an ‘OFF’ state, and an enhanced capacity to generate phenotypically heterogeneous subpopulations.

In summary, our work characterizes heterogeneity in SPI-2 expression at the mechanistic and information processing level. We show probabilistic, temporally-based tuning of the system as a way to circumvent environmental uncertainty, demonstrating that virulence regulons present rich opportunities to identify evolutionary strategies for navigating complex environmental transitions.

## ACKNOWLEDGEMENTS

We thank past and present members of the Lane Lab for their sharing of protocols and code, and valuable feedback on this manuscript at every stage. In particular we thank Cece Stumpf for her assistance with knockout strain validation and ssrB construct optimization. We also thank Chris Petersen and Elias Guan for reagents and assistance with smFISH probe labeling. We thank the Robert H. Lurie Comprehensive Cancer Center of Northwestern University for the use of the Flow Cytometry Core Facility and the Keck Biophysics Facility, which provided training and equipment for flow cytometry experiments and HPLC purification of smFISH probes respectively. The Lurie Cancer Center is supported in part by an NCI Cancer Center Support Grant #P30 CA060553. MRS was supported by the Cellular and Molecular Basis of Disease Training Program (NIH T32 GM008061).

## AUTHOR CONTRIBUTIONS

Conceptualization: MS, KL; Investigation: MS; Formal analysis: MS, KL; Visualization: MS, KL; Writing - original draft: MS, KL; Writing - review and editing: MS, KL; Funding acquisition: MS, KL; Supervision: KL.

## METHODS

### Data and Materials Availability

Flow cytometry summary statistics and quantification of imaging data are available on GitHub (https://github.com/LaneLab/Spratt_SPI2_tuning_paper). Plasmids and STm strains will be shared upon request.

### *Salmonella* Strains and Growth Conditions

*Salmonella enterica* serovar Typhimurium (ATCC strain 14028s) was a gift from the lab of Michael McClelland and was used as the WT strain in all experiments. Knockout strains including Δ*ssrA,* Δ*ssrB*, Δ*phoP,* Δ*fur,* and Δ*ptsN* were from the single-gene mutation library (Porwollik et al., 2014). Knockout strains were PCR confirmed.

For overnight cultures and uninduced control cultures, bacteria were cultured in M9 minimal medium (1X M9 Salts with 0.1mM CaCl_2_, 2mM MgSO_4_, 0.4% glucose) and supplemented with 0.2% casamino acids (M9/Glc/CAA). To induce SPI-2, bacteria were grown in pH 5.0 MgM-MES (170mM 2-[N-mopholino] ethane-sulphonic acid (MES) adjusted to pH 5.0 with NaOH, 5mM KCl, 7.5 mM (NH_4_)_2_SO4, 0.5mM K_2_SO_4_, 1mM KH_2_PO_4_, 8µM MgCl_2_) supplemented with 38mM glycerol and 0.1% casamino acids (Beuzón et al., 1999; Yu et al., 2004). Additional MgM-MES variations were made by adjusting the pH of the MES buffer using NaOH with the exception of pH 7 variations, in which MOPS buffer was used at the same concentration. Other MgM-MES variations were prepared with varied carbon sources, MgCl_2_, and casamino acids to indicated concentrations.

For all experiments, strains were streaked from glycerol stocks onto LB plates with appropriate selective antibiotics and incubated overnight at 37°C. Plates were discarded within 1 week.

### Plasmids, Cloning and Strain Generation

A complete list of plasmids generated in this study is included in Table S2. Plasmids were assembled by adopting the EcoFlex library for Golden Gate Assembly (Figure S7) (S. J. Moore et al., 2016). Custom components, including the *ssaG* and *ssaB* promoters and the *ssrB* gene, were PCR amplified with overhangs to generate level 0 golden gate vectors.

Promoter regions from STm genomic DNA were amplified using PrimeSTAR Max DNA Polymerase. The ssaG promoter contains 246 bp directly upstream of the ssaG coding region. The ssaB promoter contains 300 bp upstream of the ssaB coding region. Level 0 plasmids were generated using Gibson assembly and confirmed via Sanger sequencing. Next, level 1 single transcription unit reporter plasmids (promoter, ribosome binding site (RBS), fluorescent protein (FP), terminator) were assembled into level 1 plasmids via Golden Gate cloning. When required, transcription units were further combined into level 2 plasmids via Golden Gate. For all cloning steps, *E. coli* TG1 cells were used.

To create *ssrB* expression constructs, the open reading frame of the *ssrB* gene was amplified from the STm 14028s genome to generate a level 0 plasmid via Gibson Assembly. Level 1 *ssrB* constructs were then generated using plasmids from the Ecoflex library as described above. Tagged *ssrB* constructs were generated via Gibson assembly. The PAmCherry sequence (Shetty & Kenney, 2023) was ordered on a custom geneblock from IDT before being Gibson assembled into the plasmid. As we were using this PAmCherry sequence only as a globular domain to separate SsrB from charged tags, we also made a single amino acid substitution reported to eliminate fluorescent signal in this protein (Q109A) (Subach et al., 2009).

To prepare electrocompetent STm strains, a glycerol/mannitol step centrifugation protocol as described in (Warren, 2011) was used. Briefly, a culture of the given strain was grown to exponential phase, then pelleted at 2000 g at 4°C for 15 minutes and resuspended in cold water. A solution of 1.5% mannitol and 20% glycerol solution was then slowly dispensed into the cell suspension. Cells were then pelleted with gradual acceleration and deceleration at 2000g for 15 minutes. Supernatant was aspirated and cells were resuspended in glycerol/mannitol solution between 250 and 500x their concentration in the original culture. Following competent cell preparation, 100 ng of plasmid was transformed into the strain via electroporation at 1800 kV, 200 Ω, 25 µF. To generate strains containing two plasmids, competent cells were made of the strain containing one plasmid and then transformed with the second plasmid.

To make glycerol stocks of strains used in this paper, 700µL of an overnight culture of the strain grown in LB with required antibiotics was added to 300uL of 50% w/v glycerol, vortexed lightly and frozen at -80°C.

### *In vitro* Induction of SPI-2

STm cultures were inoculated from single colonies and cultured shaking for 14-16 hours at 37°C in 3 mL of M9/Glc/CAA with appropriate antibiotics. Cultures were then back diluted 1:100 into 3 mL of fresh M9/Glc/CAA and grown shaking for four hours. Cultures were then back diluted once more to an OD600 of 0.02 into 3-5 mL of experimental media. At indicated fixation time points, 600 µL of culture was added to 200 µL of 16% paraformaldehyde (PFA). This mixture was inverted to mix and incubated at room temperature for 30 minutes. Samples were washed twice in 1 mL of PBS and then stored in 1 mL of PBS at 4°C until flow cytometry or imaging was performed. Samples were analyzed within one week of fixation.

### Flow Cytometry and Analysis

Fixed cells were diluted into PBS such that they could be analyzed at approximately 10,000 events per second or below. Samples were analyzed using a BD LSRFortessa SORP Cell Analyzer with HTS (6-laser 18-parameter). At least 50,000 mRuby2 positive events per sample were captured.

Analysis and plotting of flow histograms was done using FlowJo version 10. Gating strategy is illustrated in Figure S8. Gates were drawn for each experiment and are consistent across samples within an experiment. Because of the presence of the mRuby2 signal in all reporter samples, light scatter gates were drawn liberally to encompass the bulk of the densest region of events. To eliminate debris events, mRuby2 positive samples were then identified by drawing a gate such that less than 0.05% of the negative control sample was classified as mRuby2 positive. mRuby2 positive events were then down-sampled to exactly 50,000 events using the FlowJo plugin DownSample (ver. 3.3.1). All further analyses and plotting were performed on down-sampled, mRuby2 positive events. Statistics including mean intensities and % GFP(+) cells were exported from FlowJo and plotted using custom Python code using Seaborn (version 0.13.1) and Matplotlib (version 3.8) packages.

### Growth Curves

Cells were back diluted from overnight cultures 1:100 into 3 mL of M9/Glc/CAA and grown in culture tubes for four hours. After four hours of growth, cells were diluted to an OD of 0.005 in 1mL of the media of interest with appropriate selective antibiotics. 150 µL of each dilution was transferred to each of 3 wells of a glass bottom Tecan 96 well plate. Well plates were sealed with adhesive plastic film, and holes were made in the top of each well with a 30 gauge hypodermic needle to allow for aeration. Cells were grown in a Tecan Spark Microplate reader. Following a settling time of 15 minutes to equilibrate temperature, OD600 was recorded every 5 minutes for 18 hours, with orbital rotation speed of 240 rpm. Growth curve data was plotted and analyzed using the Omniplate package (Montaño-Gutierrez et al., 2022).

### Fluorescence Microscopy

Images were acquired using a Nikon Ti2-E microscope with Lumencor Spectra III light engine, Prime BSI CMOS camera, motorized stage, perfect focus system controlled by Nikon Elements software and Oko environmental enclosure with temperature control. All images were captured with a 100x/1.45 numerical aperture objective with 2x2 binning and 16-bit depth.

All LED excitation intensities were used at 50% of the maximum power. Images for reporter strains after fixation were acquired on agar pads as described below at 500 ms in the Phase channel, 30 ms in GFP, and 25 ms in TRITC. Exposure times for other experiment types are included in those sections.

### smFISH

The protocol for smFISH was adapted from (Skinner et al., 2013) with adjustments to optimize the protocol for use in STm. We selected the largest gene in our operon of interest (*ssaL*) to ensure the minimum number of probes for optimal signal could be used. Probes were designed with the LGC Biosearch Technologies’ Stellaris^®^ RNA FISH Probe Designer. Probe sets were labeled using the protocol from (Gáspár et al., 2018). Briefly, oligos were purchased from IDT with standard purification. Oligonucleotide sequences are listed in Table S2. Probes were labeled with ATTO633 using a Terminal Deoxytidyl Terminase reaction and ethanol purified. Labeled probes were further purified via HPLC using an Agilent C18 Eclipse Plus column. Peak fractions were pooled and concentrated using a SpeedVac and final concentration was measured using a NanoDrop.

For sample collections, a volume of cells equivalent to 1 mL at an OD of 0.75-2.0 was fixed by adding 16% PFA to a final concentration of 4%, inverting to mix, and incubating at room temperature for 30 minutes. Samples were washed twice in 1mL PBS and then resuspended in 300µL of DEPC-treated water. To permeabilize cells, 700µL of 100% ethanol was added to the resuspended cells for a final concentration of 70% and incubated overnight at 4°C. Following ethanol permeabilization, cells were washed once in PBS and resuspended in 1 mL of lysozyme buffer (100µL Tris-HCl pH 8, 100 µL EDTA pH 8, 100 µL of 10X PBS, 700 µL of DEPC-treated water) with 100 µg/mL of lysozyme. Samples were incubated rocking at room temperature for 45 minutes. All wash steps following permeabilization were performed at 1000 g for 5 minutes. Cells were washed once in 1 mL of formamide wash buffer (100 µL 20X SSC, 353 µL formamide, 547 µL DEPC-treated water) and resuspended in 50 µL of hybridization buffer (prepared as described in Skinner S. et al.) with 2.66 nM/probe used per reaction. Samples were hybridized rotating at 30°C protected from light for 16-18 hours.

Following probe hybridization, 200µL of wash buffer was added to samples. Samples were centrifuged and cells were resuspended in 200 µL of wash buffer and incubated rotating at 30°C for 30 minutes. Samples were centrifuged and this wash step was repeated once more. Following washing, cells were resuspended in 2x SSC buffer prior to imaging. Samples were stored protected from light at 4°C and imaged on agar pads within 5 days of hybridization. Images for smFISH acquisition were captured at 400 ms exposure for Cy5, 50 ms exposure for GFP, and 500 ms exposure for Phase.

### Activation and Growth Simulations

To simulate changes in SPI-2(+) and SPI-2(-) populations to distinguish between ‘Founder’ and ‘Time-Delayed’ hypotheses, we simulated the populations with a simple probabilistic simulation. Simulations were set over 361 time steps, and the starting cell population was set at 2x10^7^ cells (approximately the number of cells in a dilution to an OD of 0.02 as in our experiments). At every time step, we used a fixed probability of 0.005 to randomly define how many cells transitioned into a SPI-2 ON state, and varied growth rates for the SPI-2(+) and SPI- 2(-) populations. We used an average doubling time of 240 minutes (approximately the rate we observe in culture (Figure S1.1)). We varied the doubling time down to 80 minutes, which would be closer to the rate expected in rich, pH optimal media. We also varied the time window at which transitions were able to occur. This simulation assumes that there is no ON to OFF switching, and further assumes that progeny of a SPI-2(+) cell are also SPI-2(+). For each tested scenario, we extracted the number of SPI-2(+) and SPI-2(-) cells across time.

### Immunofluorescence

For immunofluorescence of HA-tagged proteins, STm cultures were prepared as in the *in vitro* induction of SPI-2 protocol. A sample was fixed after four hours of growth in the first back dilution in M9/Glc/CAA and after two hours of growth in the second back dilution in MgM-MES in a volume roughly equal to 1mL at an OD of 0.5-1.0. Fixed cells were washed twice and then permeabilized as described in the smFISH protocol.

Following permeabilization, samples were incubated rotating in blocking buffer consisting of PBS supplemented with 5% goat serum, 0.05% Tween and 30 mg/mL BSA for 30 minutes. After blocking, cells were resuspended in 100 µL of blocking buffer and 1:200 dilution of rabbit monoclonal anti-HA antibody. Cells were incubated rotating with primary antibody for 1 hour. Cells were washed 3x with 1 mL PBS, then resuspended in 100 µL of blocking buffer and 1:1000 secondary goat anti rabbit Cy5 antibody. Cells were incubated rotating protected from light for 1 hour. Cells were washed 3x with 1 mL PBS with 5 minute rotations at each wash. Cells were spotted and imaged on agar pads. All steps were carried out at room temperature, unless noted. Images for immunofluorescence were captured at 500 ms exposure in the Phase channel, 100 ms in Cy5, 70 ms (3xHA:sfGFP experiments) or 20 ms (*ssrB* with reporter experiments) in GFP, and 20 ms in TRITC (*ssrB* with reporter experiments only).

### Agar Pad Imaging of Fixed Cells

Agar pads were prepared by melting 2% low melt agarose in PBS. Approximately 2 mL of melted agarose was pipetted onto an ethanol cleaned coverslip. A second coverslip was then set on top of the melted agarose. This agar pad ‘sandwich’ was allowed to dry for at least 45 minutes. Following solidification, the coverslips were removed and the pad was cut up into roughly 1 cm squares. Cells were diluted to image with sufficient cell separation to avoid segmentation issues. 1-2 µL of diluted bacterial sample was pipetted directly onto the agar pad and allowed to dry for 5 minutes, protected from light if necessary. The pad was then flipped onto a fresh ethanol cleaned cover slip and imaged within an hour of agar pad preparation.

### Dual Input Mother Machine (DIMM) Preparation, Loading, and Imaging

A silanized DIMM master mold was obtained from Micro Resist Technology GmbH (Kaiser et al., 2018). PDMS was combined at a ratio of 1:10 curing agent to PDMS base, degassed, and poured over the master. The devices were incubated overnight at 65°C, then removed from the master and cut. Tubing holes were punched with 0.75 mm biopsy punches. Devices were plasma cleaned using a Harrick Plasma Cleaner on high for 40 seconds, then incubated at 65°C for 15 minutes to 1 hour. Devices were bonded at least 24 hours in advance of an experiment.

On the day of the experiment, devices were passivated by injecting passivation buffer (denatured Salmon Sperm DNA and 10 mg/mL BSA in a 1:3 ratio) through the cell inlet while simultaneously injecting water through the waste outlet. Passivated devices were incubated at 37°C for 1 hour. Following passivation, MgM-MES and M9/Glc/CAA supplemented with 2g/L Pluronic and 1% passivation buffer were injected through the device for 30 minutes to an hour at 1.5µL/min using Harvard Apparatus Pump 11 Elite syringe pumps.

Prior to passivation, cells were back diluted from overnight cultures and grown for 4 hours in M9/Glc/CAA with 2g/L Pluronic. Cells were concentrated roughly 50x via centrifugation and then injected through the cell inlet with a 1mL syringe. Pressure was maintained at the cell syringe and the media syringes to stabilize flow until cell loading was evident. Flow was monitored at the dial-a-wave junction using UV fluorescent beads to ensure the media transition component of the device was functional. After loading, media was injected from both syringes at 1.5µL/min for 5 minutes or until cells were cleared from the main channel. Rates were then transitioned to 2.4µL/min for M9/Glc/CAA and 0.6µL/min for MgM-MES for at least 2 hours to allow cells to recover and grow into channels in the complete absence of inducing media. Following this recovery period, cells were imaged in M9/Glc/CAA for at least 5 hours. The flow rate was then changed such that MgM-MES was at 2.4µL/min and M9/Glc/CAA was at 0.6µL/min to expose the cells to SPI-2 inducing conditions. Flow rates for media transitions were selected based on published DIMM protocols (Kaiser et al., 2018). Cells were then grown and imaged in MgM-MES for 18 hours. Images for DIMM experiments were captured every 5 minutes, at 50ms exposure for sfGFP(LVA), 30ms exposure for mRuby2, and 500ms exposure for Phase.

### Image Analysis

#### Fixed cell reporter and immunofluorescence

For fixed cell reporter and immunofluorescence images, cell segmentation on the Phase channel was performed using Omnipose (Cutler et al., 2022) with the ‘bact_phase_omni’ pretrained model, a flow threshold of 0, and a mask threshold of 1.5. Fluorescent intensity measurements, area, eccentricity, and solidity were extracted using sci-kit image regionprops (version 0.22.0). Cell labels were filtered to remove debris on the basis of area (labels between 50 and 250 pixels retained), eccentricity (labels >= 0.6 retained), solidity (labels >= 0.8 retained), and for cells containing a reporter plasmid, mRuby2 median intensity (labels with a median intensity greater than two times the average of cells in a mRuby2 negative control were retained).

#### smFISH

Omnipose was used to segment cells on the Phase channel with identical settings as for fixed cell reporter and immunofluorescence images. Spot detection was performed using FISH-quant-v2 (version 0.6.2) (Imbert et al., 2022) after application of the built in LoG filter with a voxel size of 103x103 nm and a spot radius of 105x105 nm. Spot detection thresholds were set on a per experiment basis as the mean of the automatic thresholding function for all WT and ΔssrB strain images plus two standard deviations. Because of slight offsets between the phase and the fluorescence channels, spots found outside of cells but within a 2 pixel radius of a cell label were assigned to the cell with the greatest overlap. Debris filters were applied as described above based on area, eccentricity, and solidity. Spot intensities were extracted as the value of the spot centroid as detected by FISH-quant-v2.

#### DIMM analysis

Individual growth channels were manually cropped. We used only growth channels of one length, but varied widths (1.1-1.6 µm) as we observed varied efficiency in channel loading across the device. For experiments in which microscope drift occurred greater than the width of the standard crop size, a rigid body transformation was applied from the PyStackReg library to align the images. Omnipose was used to segment cells based on mRuby2 intensity using the ‘bact_fluor_omni’ pretrained model, a flow threshold of 0, and a mask threshold of 2. The TrackMate FIJI plugin (Ershov et al., 2022) was used to extract data on all cells in a channel and track mother cell lineages. Lineages were tracked using the ‘Overlap Tracker’ with ‘Precise’ IoU calculation, a minimum IoU of 0.2, and a scale factor of 1.3. Minor segmentation errors were corrected in Napari (Sofroniew et al., 2025). Tracking errors were manually corrected in TrackMate. Grossly missegmented cells were manually removed from the tracks during this process. Lineages that did not fully grow into the channel, crowded the channel to the extent that segmentation or tracking were not feasible, or in which the mother cell died prior to dividing were removed from the data set. Cell features, including intensity and size measurements, were extracted using TrackMate for all tracked data. For untracked cell data, intensity and size measurements were extracted using regionprops or TrackMate equivalent definitions. Data was extracted from Trackmate XML files using pyTrackMateXML and custom code and stored in Xarray format for further analysis.

To account for day to day variability in device background intensity, we measured ten 100 pixel squared background regions from five randomly selected images per experiment. We used the mean of these background measurements to adjust cell intensity measurements such that all experiments had the mean background intensity of the brightest experiment. GFP(+) threshold was set as 3x the average normalized median intensity of all cells at time 0 in the experiment. PssaG-sfGFP(LVA) median intensity was normalized across experiments as described in Figure S6.1 and smoothed with a 3 frame rolling average for all analyses. SPI-2 state transition rates and probability based on this GFP(+) definition were quantified as described in Figure S6.3.

We categorized cells into switch-like, gradual, or non-responsive trajectories on the basis of the slope of the log(PssaG-sfGFP(LVA)) signal over time (Figure S6.4). Briefly, we selected a slope threshold of 0.04 AUs/min to define the start and end of an increase in reporter signal. We defined the start as the first time this threshold was exceeded. We defined the end as the first time following the start that the threshold was not met for at least 5 frames. If the GFP(+) threshold was crossed between the detected start and end times, cells were classified as switch-like. If the GFP(+) threshold was crossed outside of this time window, cells were defined as gradual. If cells failed to reach the GFP(+) threshold, they were classified as non-responsive.

The length of cells was extracted as the ‘feret_diameter_max’ as defined by regionprops (scikit-image). The elongation rate was calculated as an average across a given cell cycle.

#### Random Forest Regressor

We created the RFR model using the RandomForestRegressor module from scikit-learn (ver. 1.3.2). We performed the training and testing on switch-like progeny cells that were born below the GFP(+) threshold, had at least 6 hours of tracking data, and had a SPI-2 response within a fully tracked cell cycle. This population was split into 70% training data (n=413 cells), 30% test data (n=178 cells). We used 100 decision trees within the model (estimators). For cycle based features both ‘Resp. Cycle’ and ‘Average’ of the feature were incorporated. ‘Resp. Cycle’ indicates the value of that feature for the cell cycle in which the response occurred. ‘Average’ indicates the mean value of that feature for all fully tracked cell cycles for that cell. Model accuracy was calculated as 100-MAPE between experimental and predicted values. The contribution of each feature to the model predictions was evaluated using the SHAP package (ver. 0.47.2). Features incorporated into our model are defined below:

#### Cycle Features

End Length - the length of the cells at the completion of each cell cycle.

Start Length - the length of the cells at the first time point of each cell cycle.

Cycle Duration - the time from start to division for a cell cycle.

Growth - the difference between start and end length for a cell cycle.

Elongation Rate - the total growth for a cycle divided by the cycle duration.

#### Other Cell Features

Maximum mRuby2 Intensity - the highest median mRuby2 intensity measured for the cell.

Mean Lifetime mRuby2 Intensity - the average median mRuby2 intensity of the cell.

Initial Mother Cycle Duration - the time from the media transition to the first division of the original mother cell that the cell originated from.

#### Experimental Setup Features

Experimental Replicate - the replicate the cell was measured in.

Channel Width - the width of the channel the cell was growing in.

---

**Figure S1.**
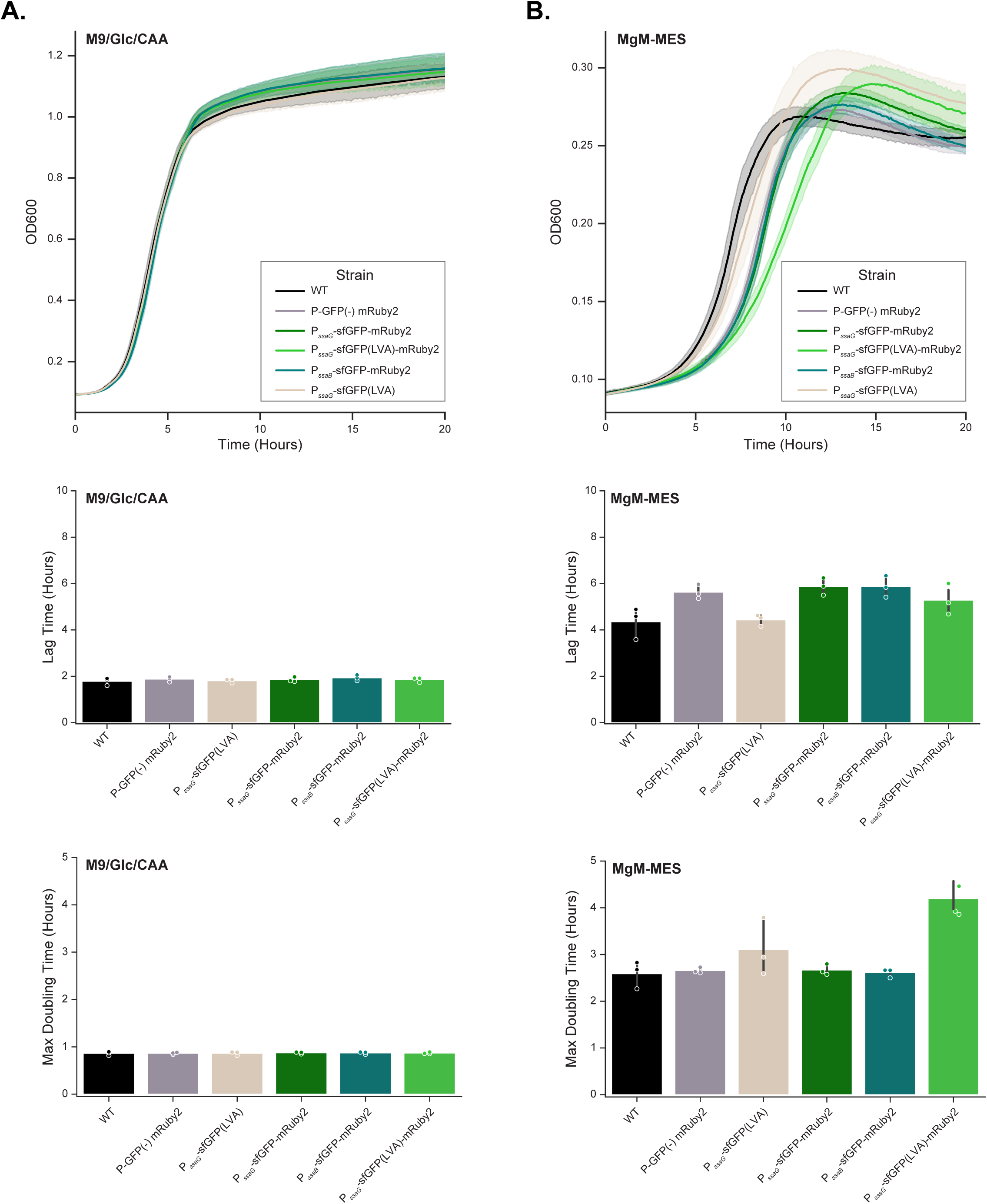
1. SPI-2 reporters cause marginal growth deficits in inducing conditions Plate reader growth curves and statistics (lag time, max doubling time) for strains grown in (A) M9/Glc/CAA or (B) MgM-MES. Data for a strain with no reporter plasmid (WT), a strain expressing mRuby2 with no promoter driving sfGFP(LVA) (Promoterless), PssaG-sfGFP with mRuby2, PssaG-sfGFP(LVA) with mRuby2, PssaB-GFP with mRuby2, and PssaG-sfGFP(LVA) with no mRuby2 are shown. Growth curves show average measurement across 3 biological replicates, each with 3 technical replicates, with error bars corresponding to standard error across the replicates. Points on statistics plots are per replicate.

**Figure S1.**
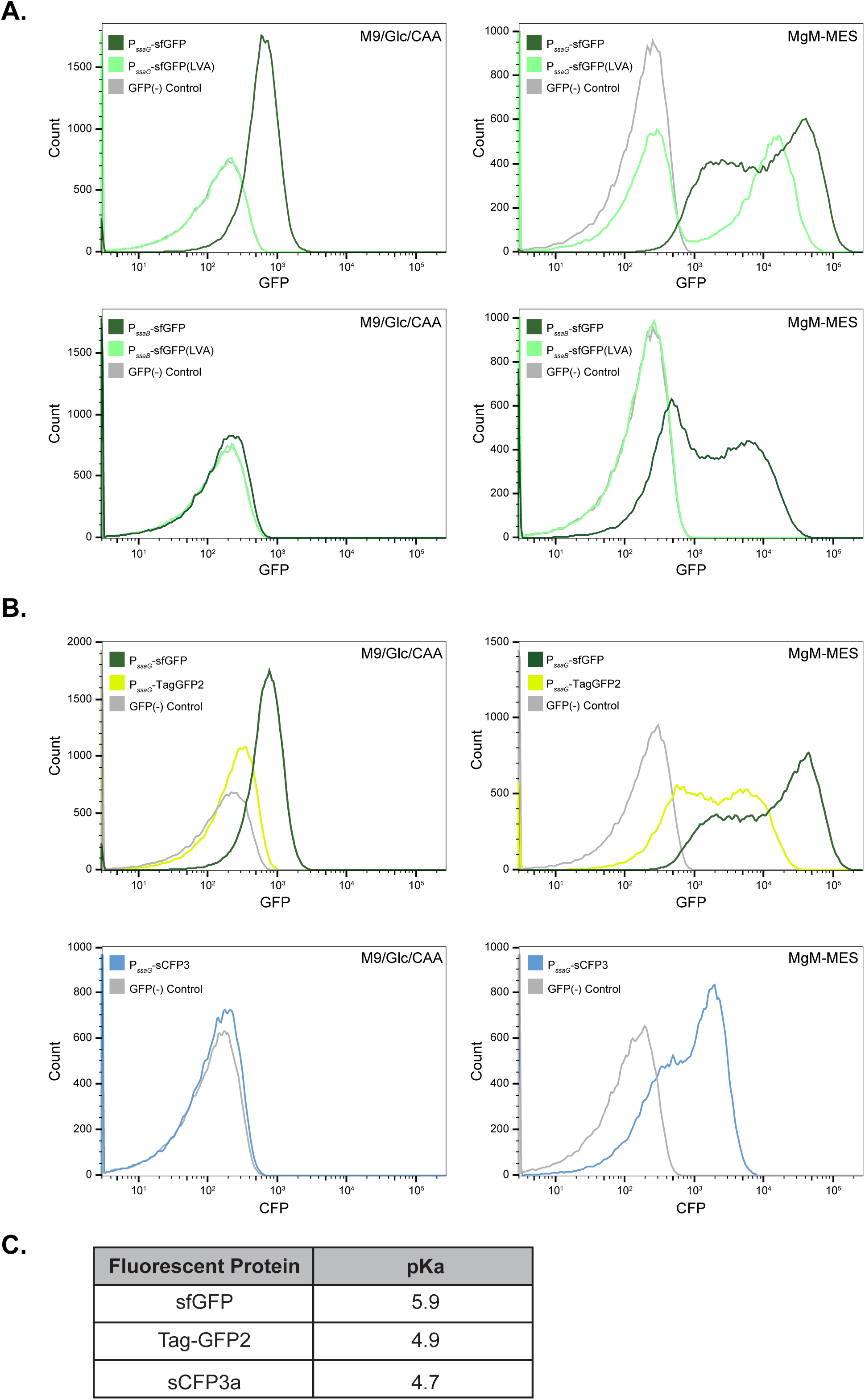
2. Optimizing the signal from SPI-2 reporter constructs (A) Evaluating the use of a LVA degradation tag: PssaG and PssaB were fused to sfGFP and sfGFP(LVA). Reporters were grown in M9/Glc/CAA and MgM-MES for 4 hours, fixed and analyzed by flow cytometry. Representative histograms are displayed, n = 50,000 cells per sample. sfGFP accumulates in the PssaG strain without a LVA tag, even in non-inducing conditions. While bimodality in PssaG expression is still observed in the absence of the LVA tag, sfGFP(LVA) improves the separation of the two subpopulations. Expression of sfGFP(LVA) from PssaB does not result in any detectable signal via flow cytometry, so sfGFP was selected. (B) Comparison of fluorescent proteins with different pKa: variations of the PssaG reporter constructed with Tag-GFP2 and sCFP3, fluorescent proteins with lower pKa values than sfGFP (C). Both fluorescent proteins show bimodal expression in inducing conditions, validating that bimodality was not an effect of sfGFP pKa. As Tag-GFP2 and sCFP3 are considerably dimmer than sfGFP, we selected sfGFP for the reporter constructs. n = 50,000 cells per sample.

**Figure S1.**
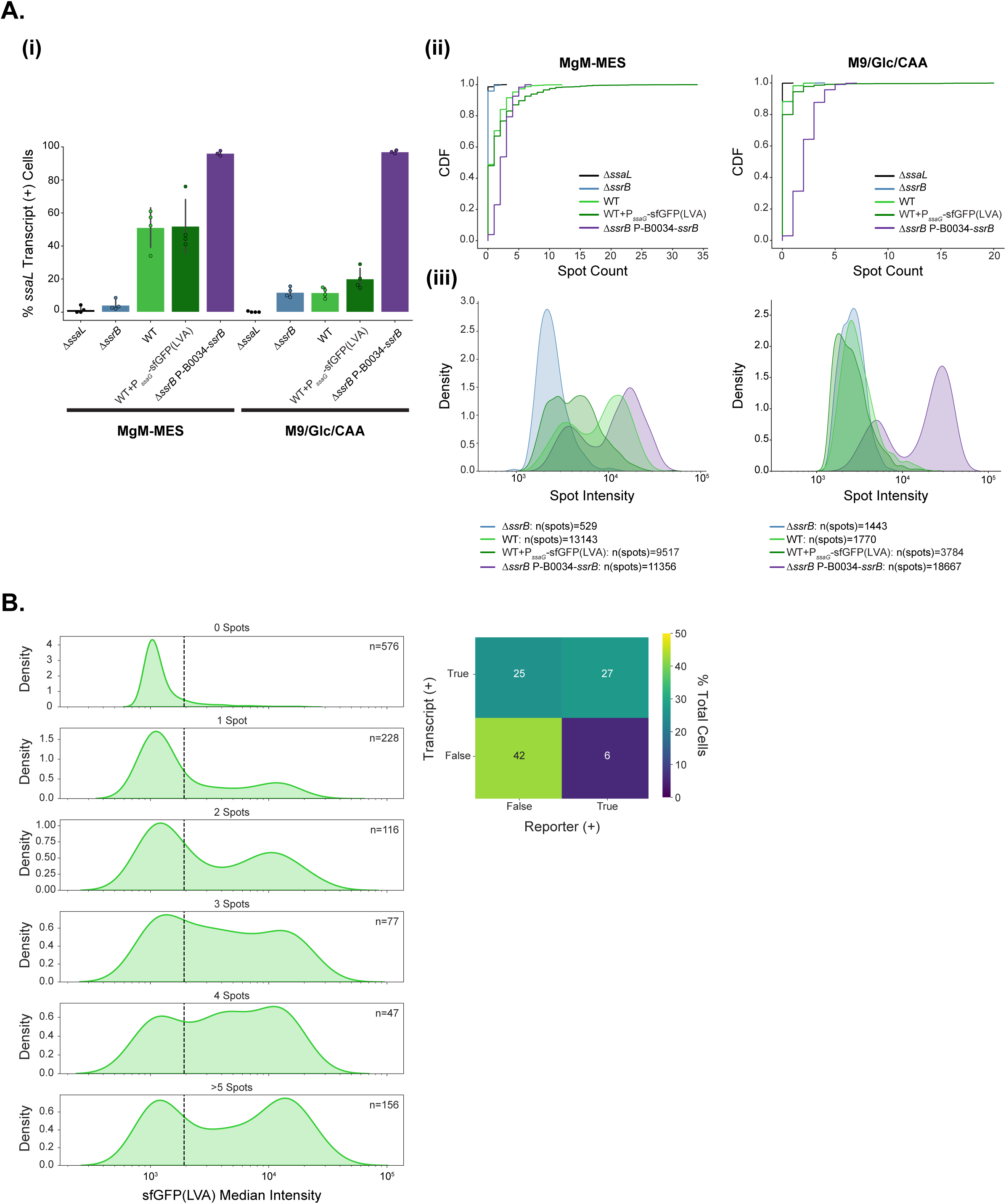
3. smFISH on ssaL transcripts shows expected activation through the canonical pathway and trends with reporter expression To evaluate if endogenous transcripts are consistent with reporter-based results during manipulations of the canonical pathway as in Figure S1.5, smFISH was performed on the ssrB KO and constitutive ssrB expression strains, as well as a strain containing the PssaG reporter alongside the WT and ssaL KO samples displayed in Figure 1C. The reporter construct used does not contain constitutive mRuby2 to avoid background signal in the smFISH Cy5 channel. (A) Percentages of cells with detectable *ssaL* transcripts are shown for each strain in inducing or non-inducing conditions (i). The detected spot count is displayed as CDFs. n=1200 cells for each strain and condition from four experiments, with the exception of the ssrB overexpression (OE) strain, which contains 900 cells from 3 experiments (ii). The distribution of spot intensity for all detected spots across all experiments is displayed for each strain, n=number of spots across all experimental replicates (iii). (B) WT+PssaG-sfGFP(LVA) cells were binned based on the number of *ssaL* spots detected, and the median intensity of the reporter per cell is shown per bin. n indicates how many cells contain the given number of spots. The black line indicates the threshold used to call cells as reporter positive - twice the mean of all cells in the WT strain across experiments. The proportions of the population of induced WT+PssaG-sfGFP(LVA) reporter cells that have transcripts detected and/or reporter signal are shown. Discrepancies are expected given the short half-lives of bacterial mRNA and the maturation time and relatively longer half-life of sfGFP(LVA).

**Figure S1.**
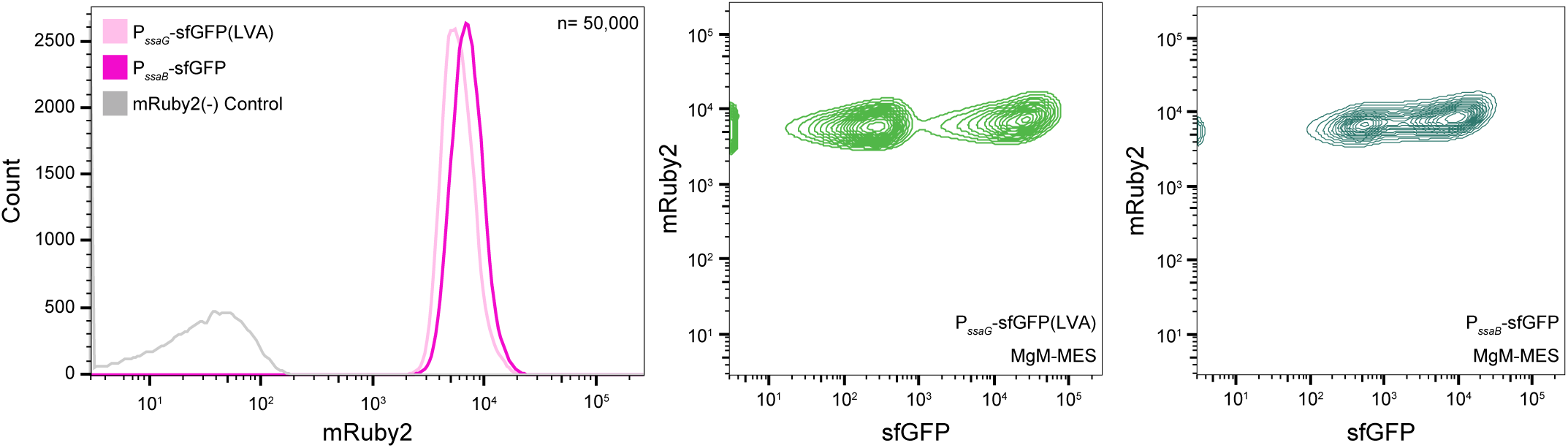
4. mRuby2 signal remains unimodal in conditions that induce SPI-2 reporter bimodality mRuby intensity from both reporter constructs after growth in MgM-MES from a representative flow cytometry experiment in Figure 1D is shown. 50,000 cells per sample are displayed. mRuby intensity versus reporter GFP intensity is shown for the same experiment.

**Figure S1.**
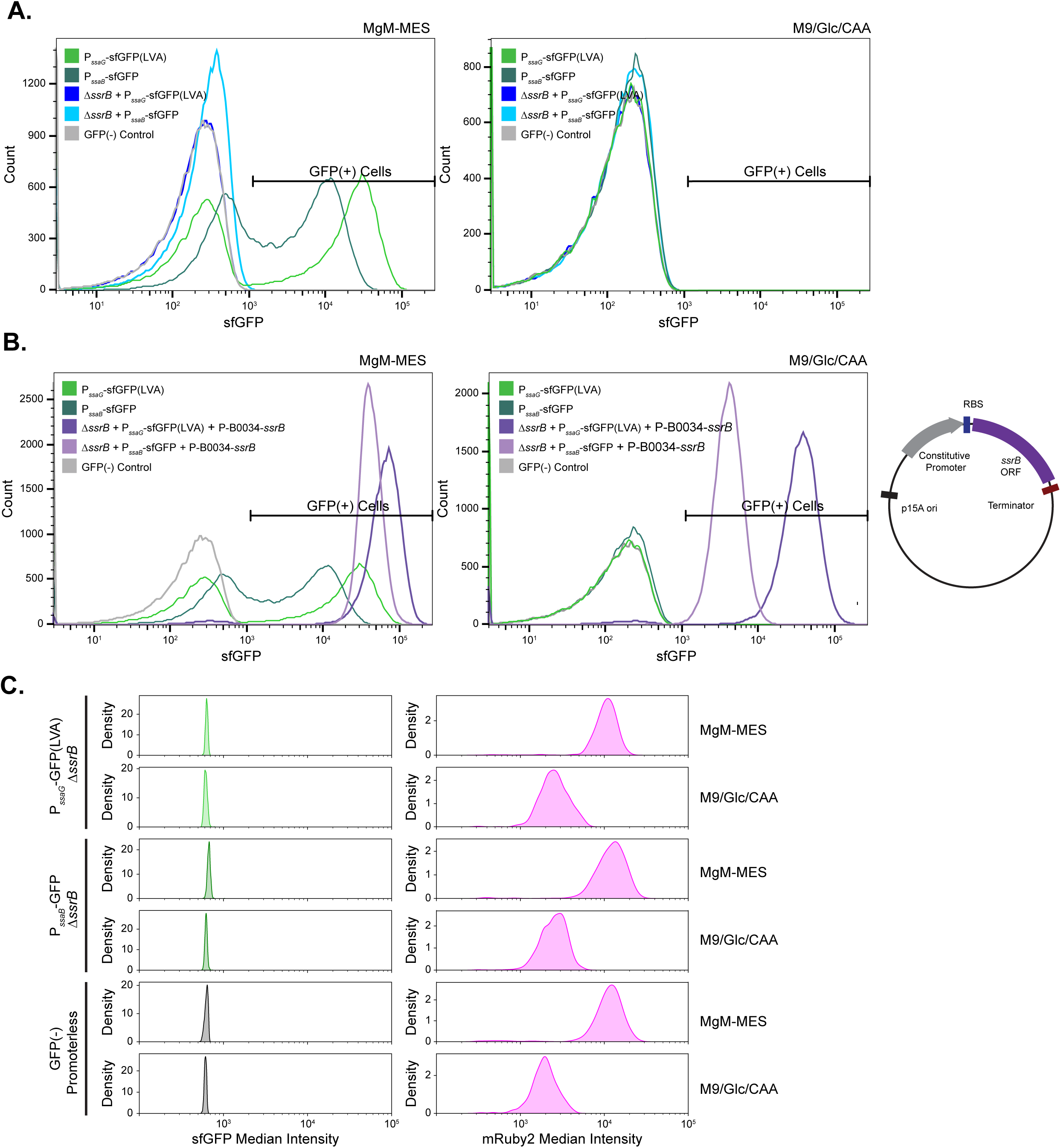
5. SPI-2 reporters are activated through the canonical SPI-2 regulatory pathway (A) WT and *ΔssrB* strains containing the SPI-2 reporters were grown in MgM-MES or M9/Glc/CAA for 4 hours, fixed and analyzed via flow cytometry. Histograms of a representative experiment are shown, displaying 50,000 cells per strain. (B) WT reporter induction was similarly compared to the *ΔssrB* strains containing a plasmid constitutively expressing *ssrB.* Right - Schematic of *ssrB* expression plasmid. (C) The SPI-2 reporters are not expressed in the absence of *ssrB*. The Δ*ssrB* strains shown in A and B alongside samples from Figure 1B were imaged and fluorescent protein expression quantified as described in the methods. Pooled data from 3 experiments are shown, n=1350 per strain, 450 cells per experiment. No GFP signal above the negative control (WT+P-sfGFP(-)mRuby2) is detected in Δ*ssrB* strains.

**Figure S2.**
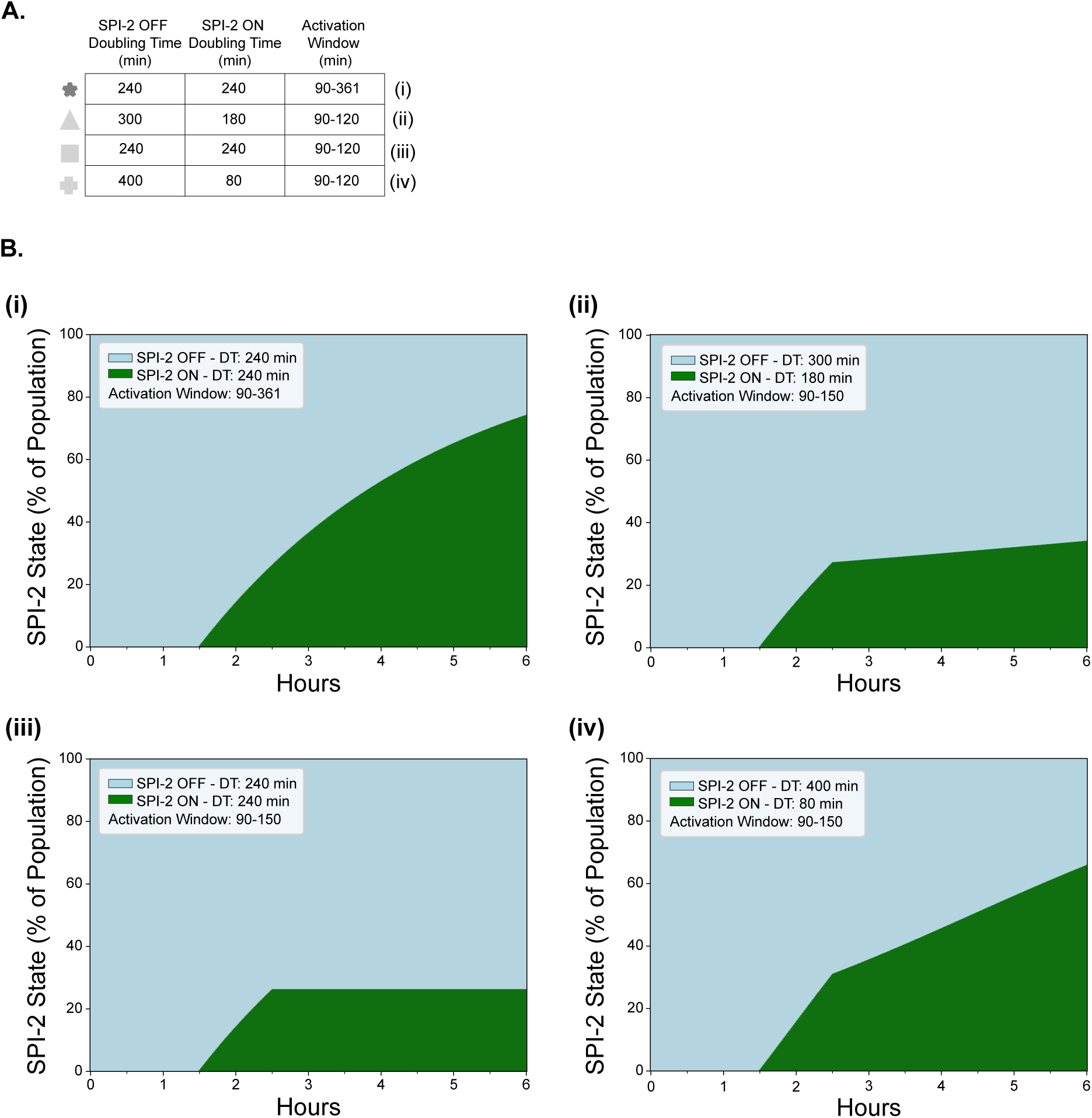
1. Founder Population vs. Time Delayed Simulations Expanded data from the simulation in Figure 2B are shown. Simulations operated as a random walk in which cells had a given probability of activation within a discrete time step (methods). For Founder model simulations, the length of time in which cells could transition from ‘OFF’ to ‘ON’ in this way was restricted (90-120 minutes), whereas in the ‘Time Delayed’ model, cells were allowed to turn on for the entire duration (90-361 minutes). (A) Simulation conditions as in Figure 2B. (B) The percentage of SPI-2 ON cells (green) and SPI-2 OFF cells (blue) is shown for each of the four simulation conditions. DT = doubling time. The increase in SPI-2(+) cells over time in the experimental data can not be explained via the Founder model without invoking an unrealistic growth advantage in the SPI-2 positive population (+).

**Figure S2.**
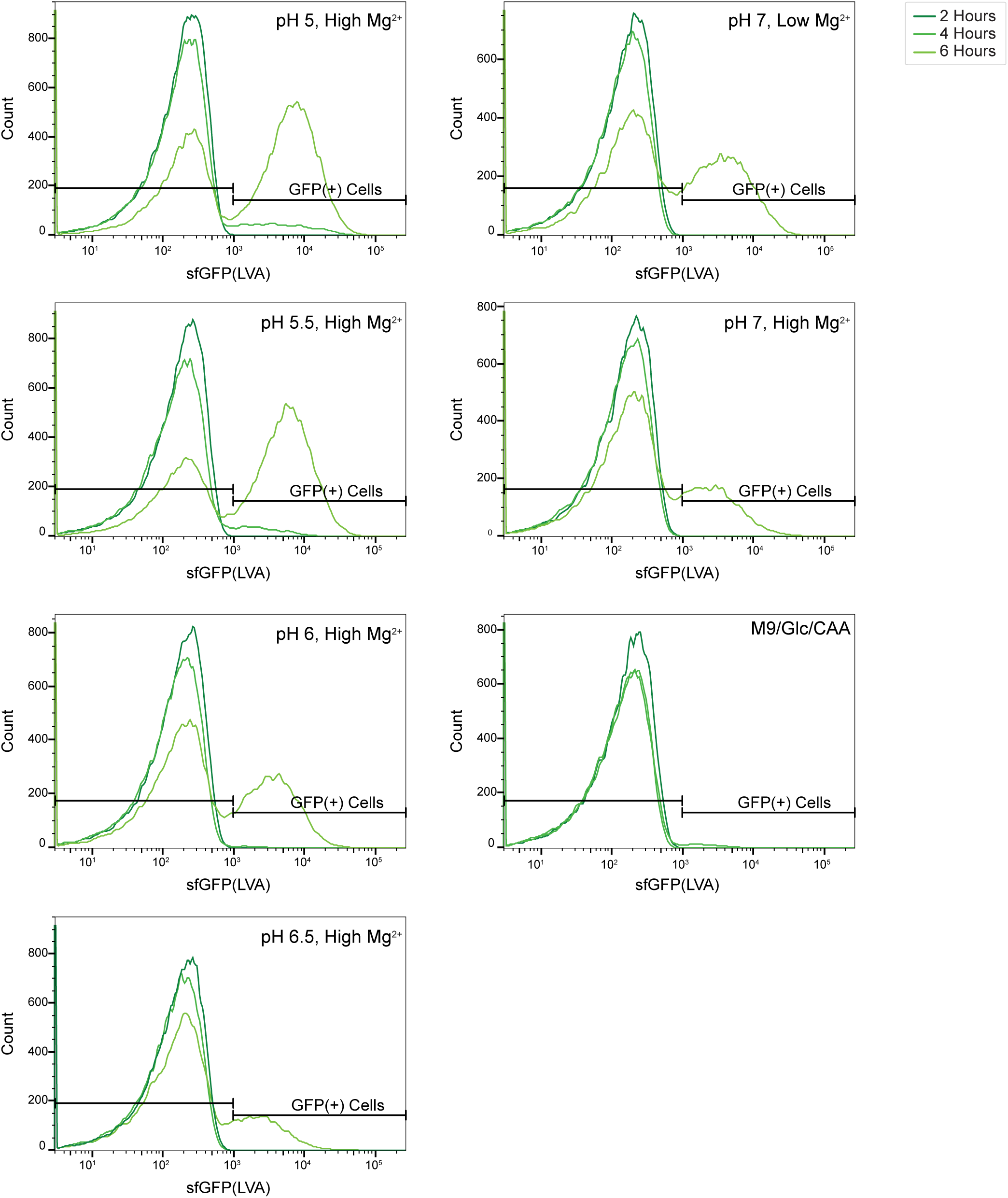
2. Bimodal SPI-2 reporter expression is maintained in varied SPI-2 inducing conditions at all timepoints Representative histograms of PssaG reporter signal across all three timepoints (2, 4, 6 hours) from a single flow cytometry experimental replicate in Figure 2C-D are displayed. Bimodality in reporter signal is maintained across time and conditions. n=50,000 cells per sample.

**Figure S2.**
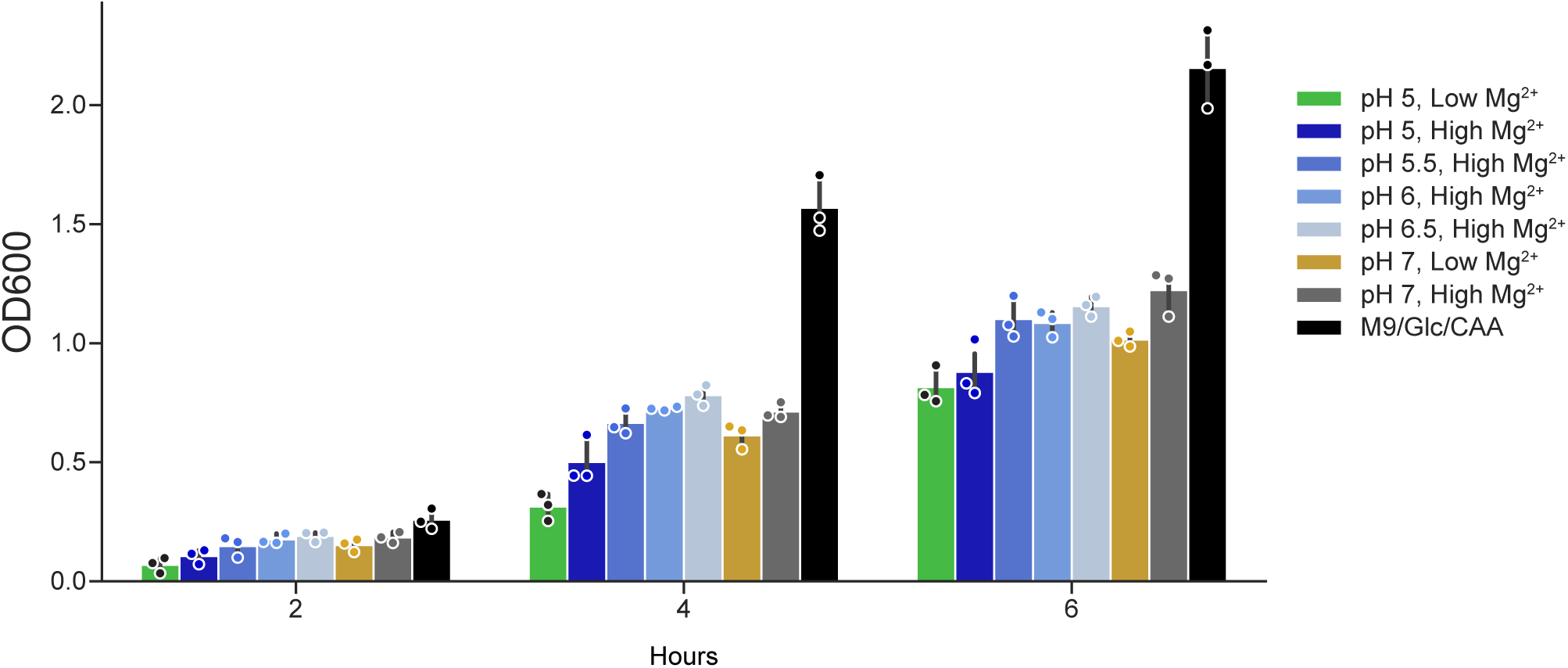
3. OD600 during growth in varied SPI-2 inducing conditions For samples in Figures 2A and 2C-D, OD600 was measured at each fixation time point. OD600 values are displayed for each condition, individual dots indicate experimental replicates. Decreased pH and minimal media conditions reduce the growth rate relative to M9/Glc/CAA. While the most growth restrictive media is standard recipe MgM-MES (pH 5, 8 µM MgCl_2_), cells divide across the entire duration in the media.

**Figure S3.**
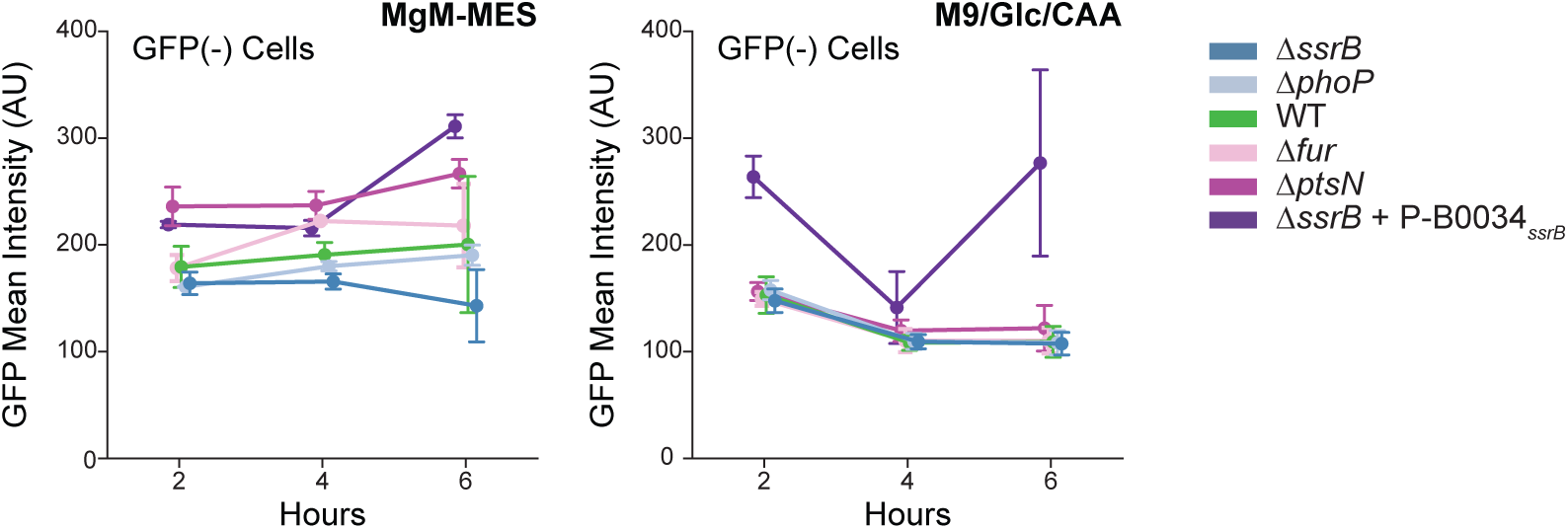
1. Mean intensity of GFP(-) populations in SPI-2 gene regulatory network knockout strains The intensity of the GFP(-) population across all samples from the 3 biological replicates in Figure 3B is displayed, error bars show standard deviation. The intensity of the GFP(-) subpopulation is maintained across timepoints and strains, consistent with single-cell transitions from SPI-2 OFF to ON rather than accumulation of the SPI-2 reporter.

**Figure S3.**
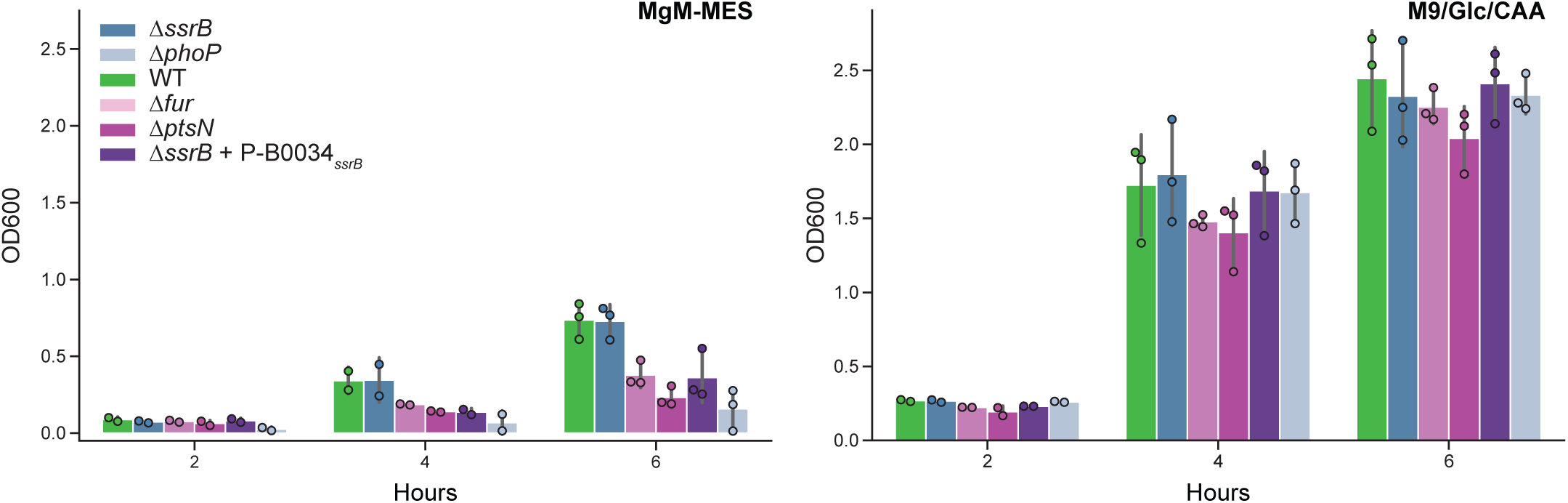
2. OD600 of SPI-2 gene regulatory network knockout strains To monitor relative growth of the different strains used in Figure 3B, OD600 was measured at each fixation time point. Average OD600 values are displayed for each strain at each time, individual dots indicate replicates.

**Figure S4.**
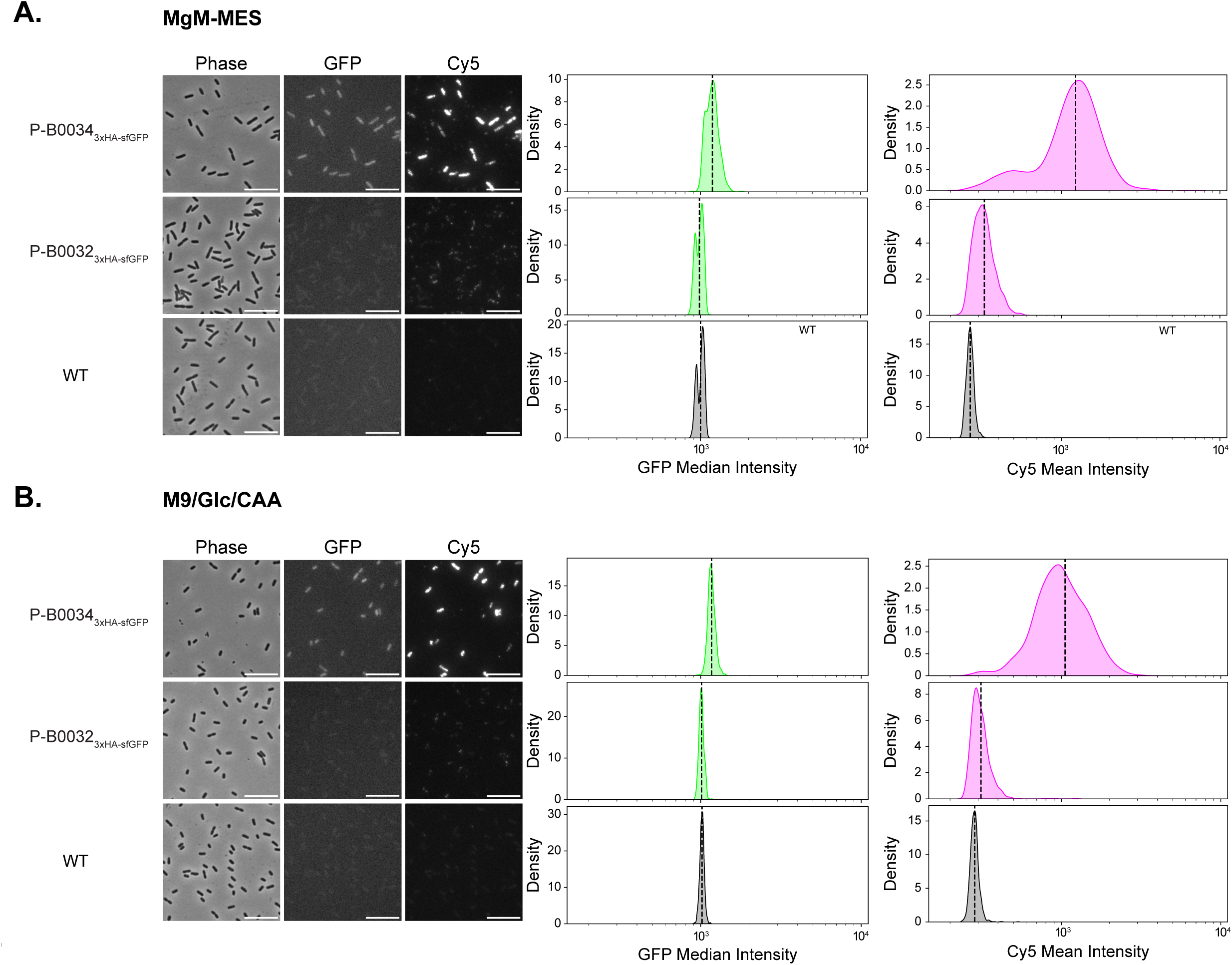
1. Synthetic RBS modulation can decrease protein production from weak expression constructs. To evaluate the ability of the varied RBS to modulate protein levels, we created constructs equivalent to P-B0032*_ssrB_* and P-B0034*_ssrB_* to express 3x:HA:sfGFP instead of *ssrB*. Representative images of the strains after 4 hours of growth in MgM-MES (A) or M9/Glc/CAA (B) followed by immunofluorescence with an anti-HA antibody are shown. Quantification of GFP intensity and Cy5 intensity (HA signal) is shown, n=1500 cells, 500 from each of 3 experiments. Mean Cy5 intensity is displayed due to the unevenly dispersed signal within the cells. Average intensity values of the populations are shown with dotted black lines. The B0032 RBS effectively reduces protein levels below the already low level of constructs with the B0034 RBS. Plasmid details are described in the methods and Table S2.

**Figure S4.**
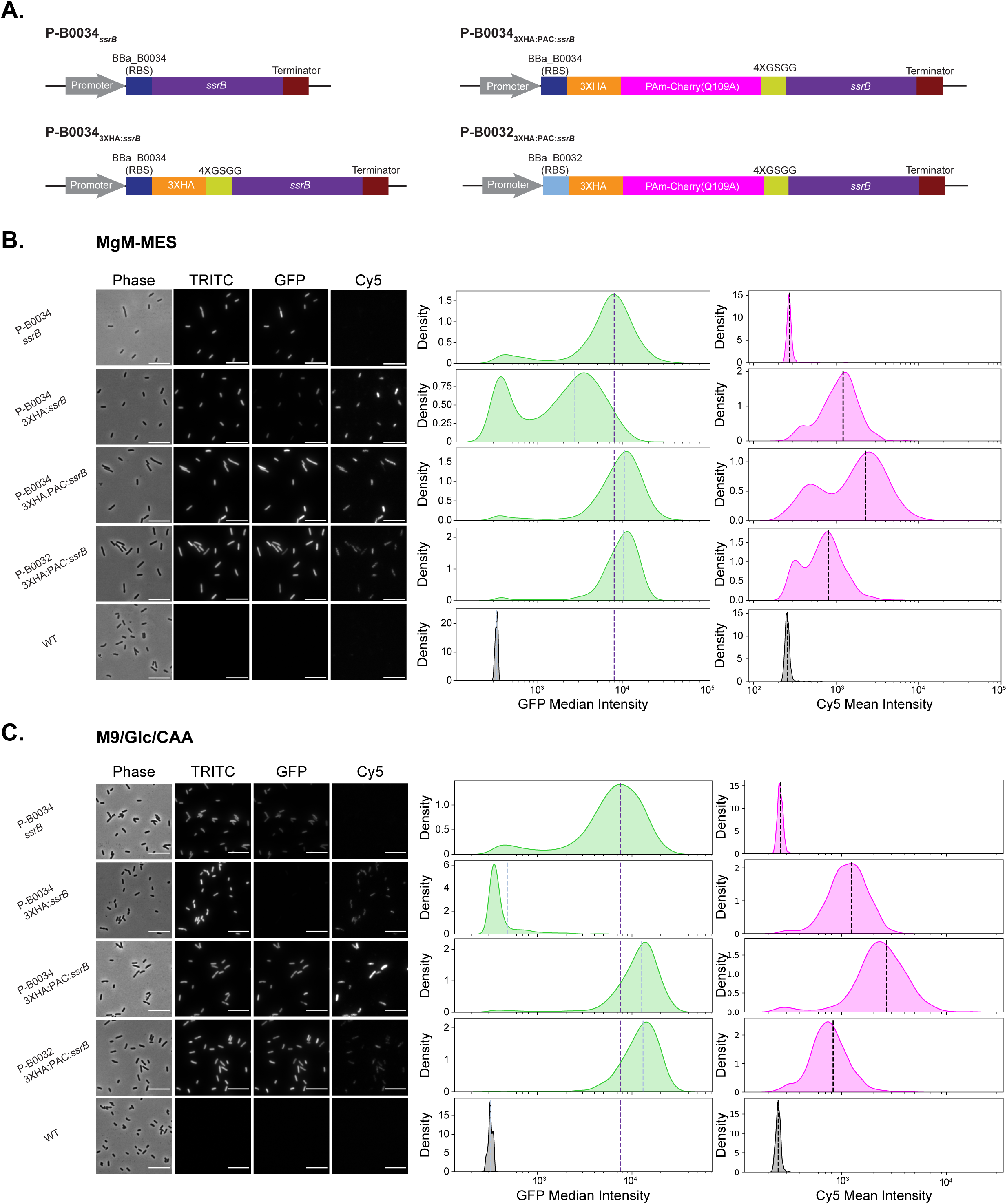
2. Tags on SsrB alter downstream gene activation Constructs with tagged versions of *ssrB* are illustrated in (A). Constructs were cotransformed with the PssaG-sfGFP(LVA) reporter into the Δ*ssrB* strain. Representative images of the strains after 4 hours of growth in MgM-MES (B) or M9/Glc/CAA (C) followed by immunofluorescence with an anti-HA antibody are shown. Quantification of GFP intensity and Cy5 intensity (HA signal) is shown, n=1500 cells, 500 from each of 3 experiments. Cells quantified were filtered based on cell size, consistent with all other imaging experiments, in this case excluding a small population of filamenting cells in some samples that may arise from the loss of the ssrB expressing plasmid in selective antibiotics. Results show that the addition of tags alters PssaG-sfGFP(LVA) expression relative to the untagged expression construct (P-B0034*_ssrB_*), with the 3XHA tag alone (P-B0034_3XHA:*ssrB*_) reducing reporter output, while the 3XHA tag with the PAmCherry spacer increases it (P-B0034_3XHA:PAC:*ssrB*_, P-B0032_3XHA:PAC:*ssrB*_). Dotted black lines show the mean of the population.

**Figure S4.**
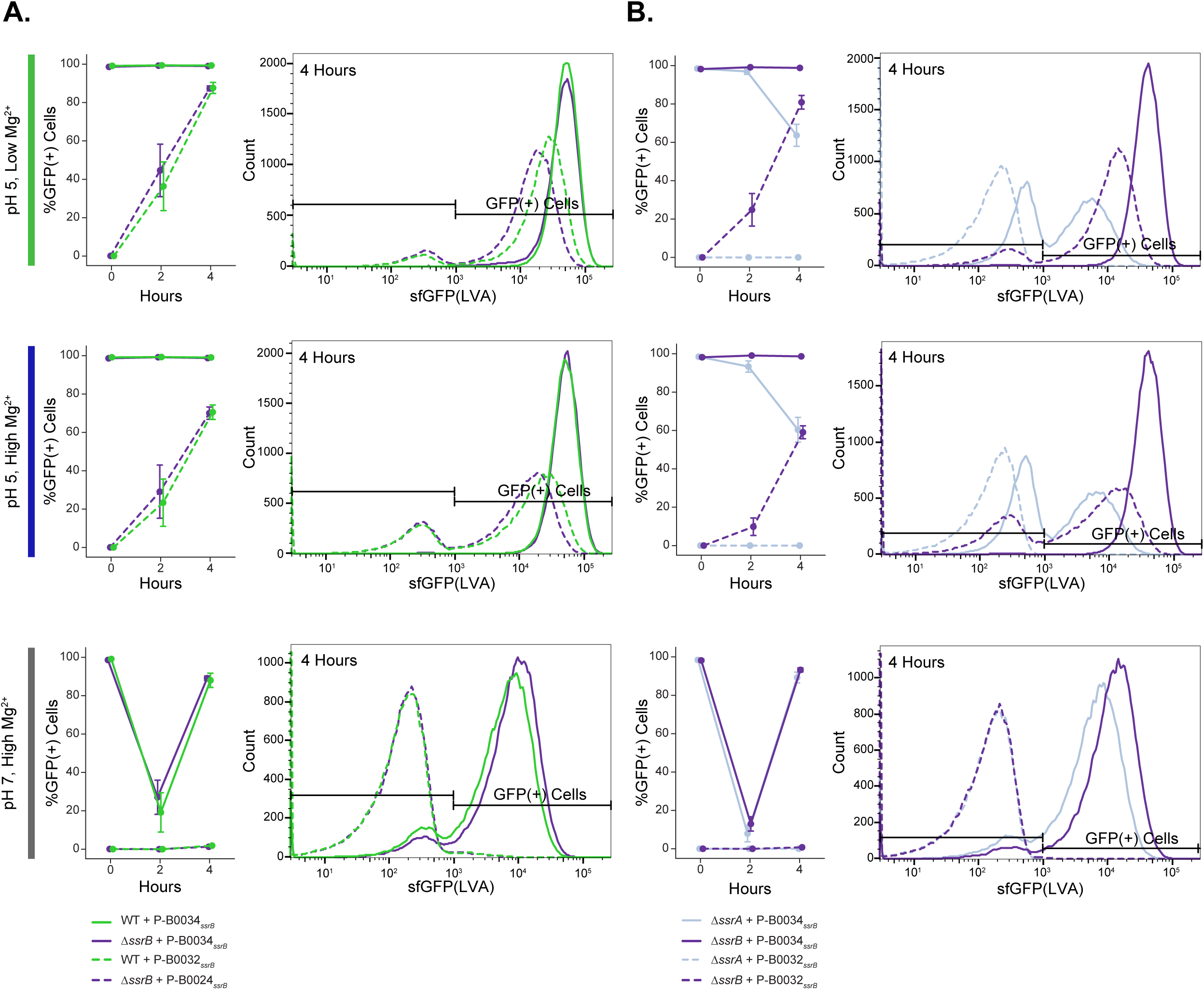
3. Reporter responses to low SsrB expression are time and environment-dependent Expanded flow cytometry data from the heatmaps in Figure 4. Percentage of GFP(+) cells across time is shown, with T=0 referring to cells grown in M9/Glc/CAA for four hours prior to dilution into respective media listed on the far left, n=3 experiments. The WT compared to the Δ*ssrB* background is shown in (A) to test the autoregulation hypothesis. The Δ*ssrB* compared to the Δ*ssrA* background is shown in (B) to test the signaling-dependent hypothesis. Representative histograms of sfGFP(LVA) intensity at four hours in each condition are displayed, n = 50,000 cells per sample. Strains are indicated below each panel.

**Figure S5.**
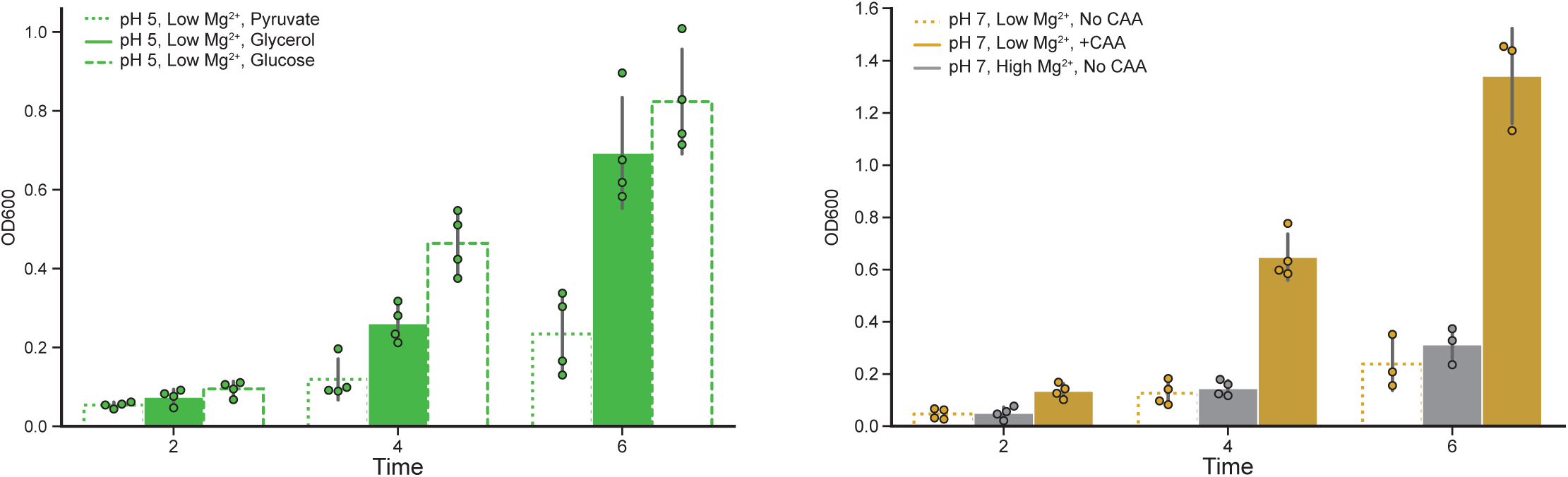
1. OD600 in nutrient variable MgM-MES experiments The OD600 readings for samples collected at the indicated timepoints in Figure 5B experiments are shown. Pyruvate slows the growth rate from the standard glycerol condition, while glucose increases it as expected. A lack of casamino acids also slows growth in weak inducing conditions relative to casamino acid supplementation.

**Figure S5.**
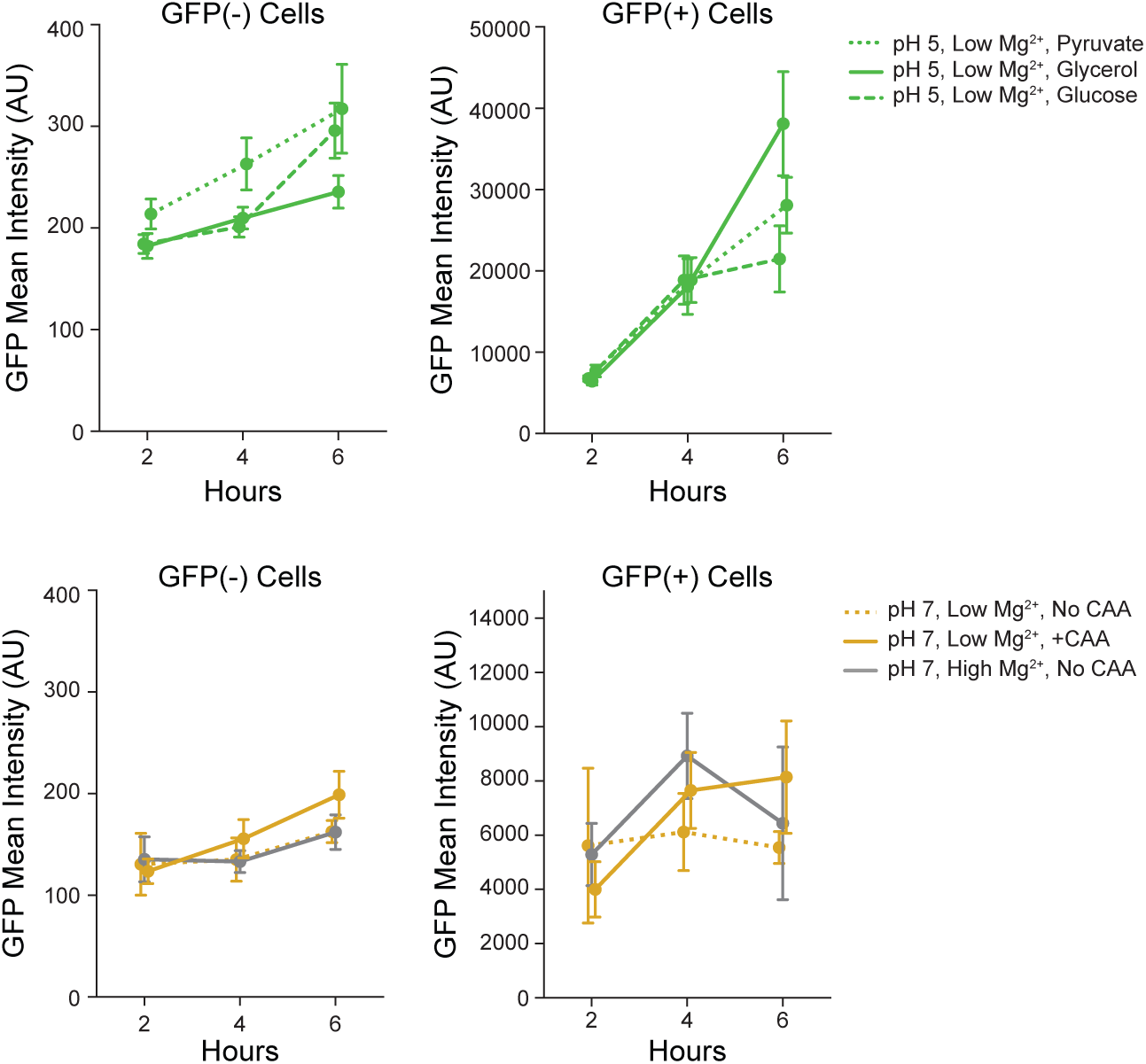
2. Mean intensity of reporter subpopulations in nutrient variable MgM-MES The mean intensities of the GFP(-) cells and GFP(+) cells are shown across time for the carbon source variable MgM-MES conditions (top) and for the CAA variable weak inducing conditions (bottom). n=4 experiments, data from Figure 5. The reporter negative populations in all conditions maintain stable, low intensities, suggesting that the ON population reflects gene expression rather than reporter accumulation.

**Figure S5.**
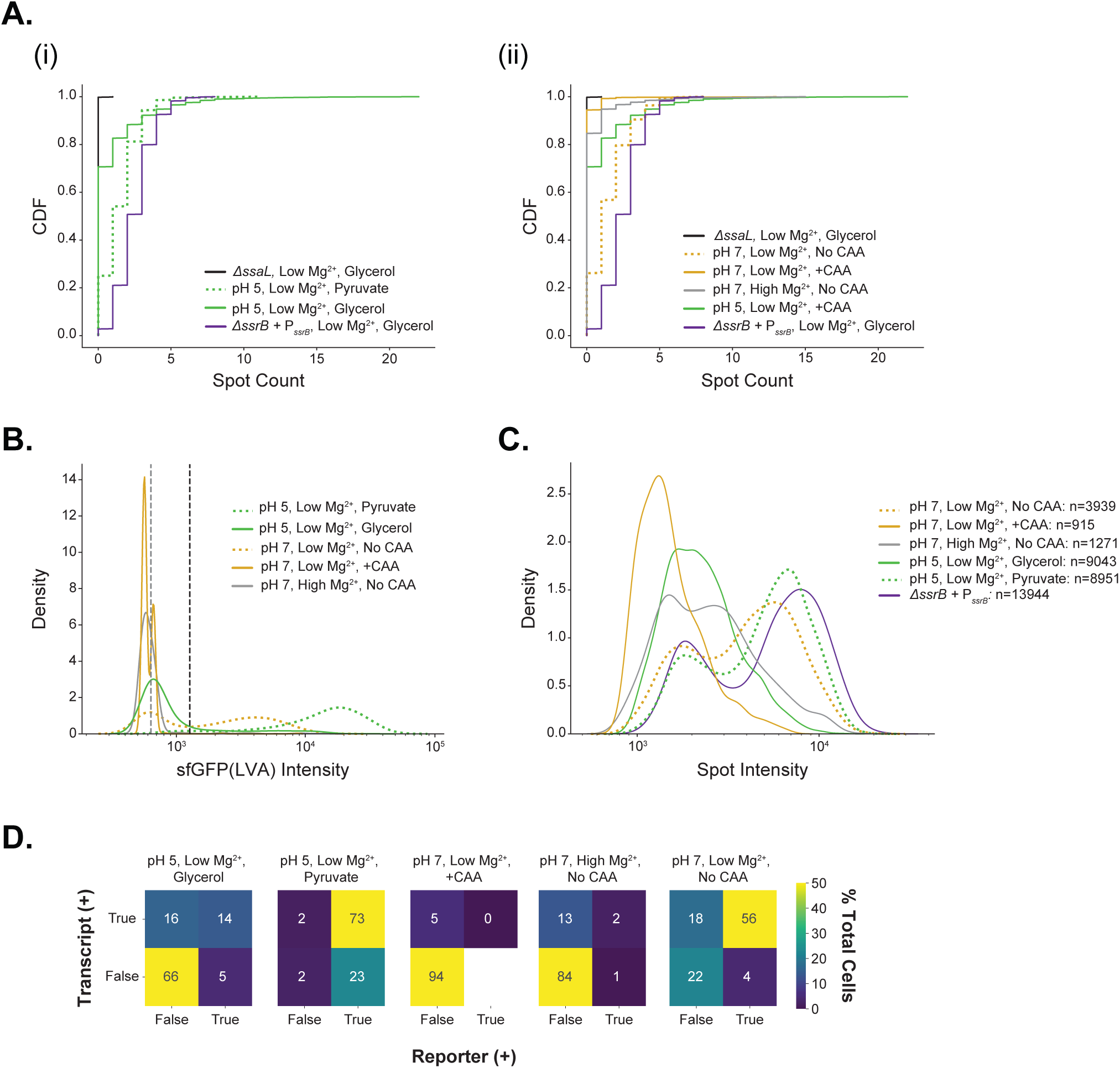
3. Reporter relative to smFISH signal in nutrient variable MgM-MES experiments Endogenous transcripts detected using smFISH are also altered by variation in growth rate through nutrient modifications, consistent with reporter data. Data is from n=900 cells across 3 independent experiments. (A) CDF of spots detected per cell are shown for the smFISH experiments in Figure 5C in which growth rate is modulated by (i) carbon source variation or (ii) casamino acid concentration, samples were collected at 4 hours of growth. (B) sfGFP(LVA) intensity distributions of cells in the same experiments containing the PssaG reporter. The mean intensity of a GFP(-) strain and the threshold used to call cells GFP(+) (double the mean of the GFP(-) strain) are shown by gray and black dashed lines, respectively. (C) The intensity per spot for each condition is shown. All spots detected within cells are included in the dataset. (D) The proportion of cells called Reporter(+), Transcript(+) (containing at least one detected *ssaL* spot), both transcript(+) and reporter (+), or negative for both SPI-2 readouts are displayed.

**Figure S6.**
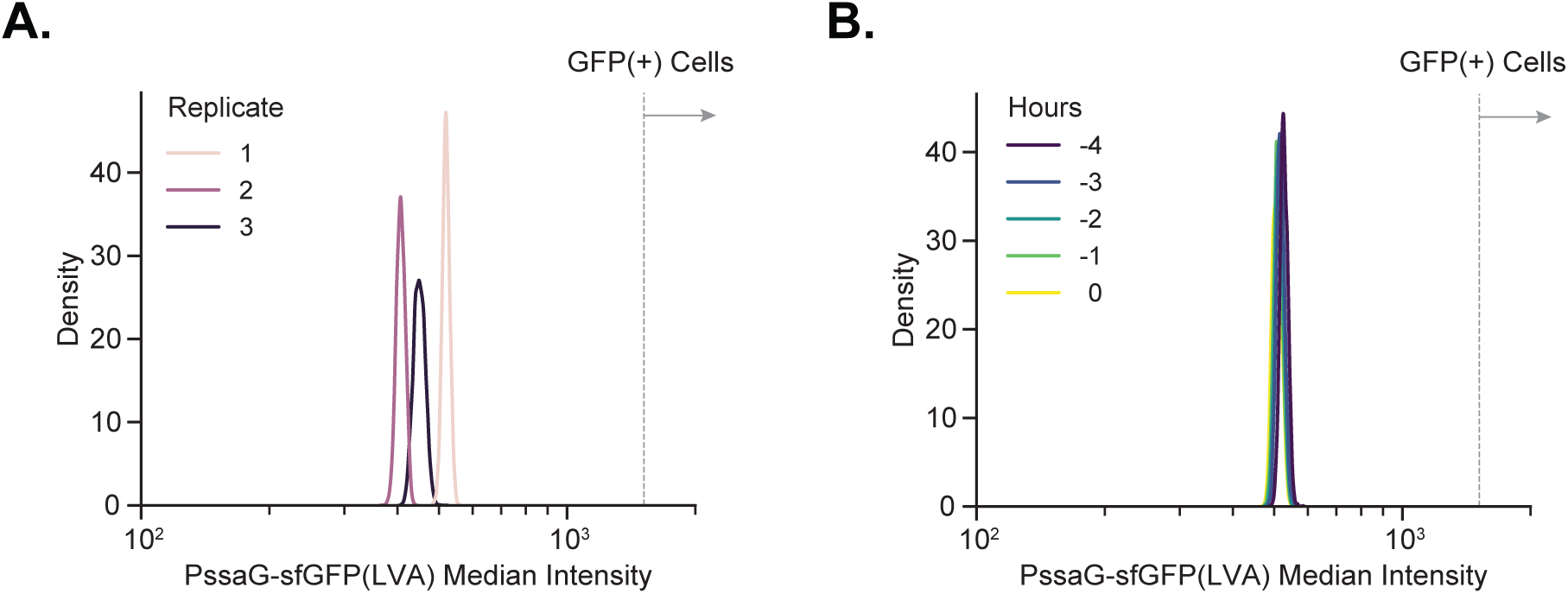
1. Reporter normalization and intensity in non-inducing media in the DIMM (A) Raw intensity values for all cells from each of three experimental replicates in non-inducing media (M9/Glc/CAA) across all timepoints. Background signal was measured for each replicate and adjusted to match the background value of the brightest replicate (#1). Adjusted values were then added to all cell intensity values. (B) Background adjusted reporter intensity pooled from three experimental replicates across time in M9/Glc/CAA shows that cells are GFP(-) before the transition into inducing media (MgM-MES). Hours are relative to the transition to MgM-MES. The dashed line represents the threshold used for calling cells as GFP(+).

**Figure S6.**
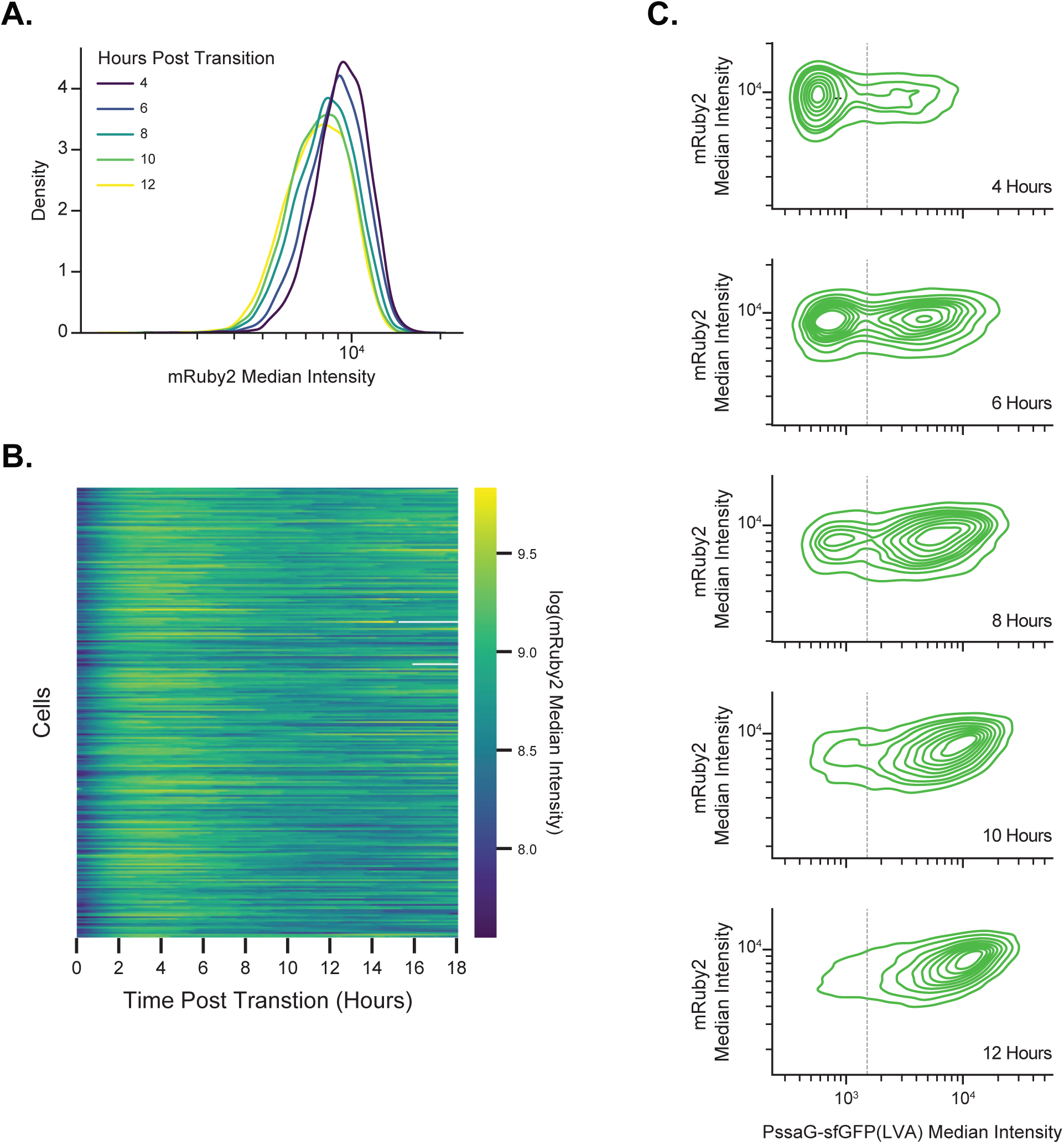
2. mRuby2 intensity is constant and uncorrelated with reporter expression during DIMM growth in MgM-MES (A) Background adjusted mRuby2 median intensity pooled from 3 experimental replicates during growth in MgM-MES, timepoints are equivalent to Figure 6B. n>4900 cells for each timepoint. (B) Mother cell mRuby2 median intensity across all timepoints. Cells are sorted based on their time to cross the GFP(+) threshold as in Figure 6D. (C) Correlation of mRuby2 intensity with PssaG-sfGFP(LVA) intensity across timepoints during growth in MgM-MES. Cells are equivalent to those in panel (A). mRuby2 expression is not correlated with PssaG reporter activation.

**Figure S6.**
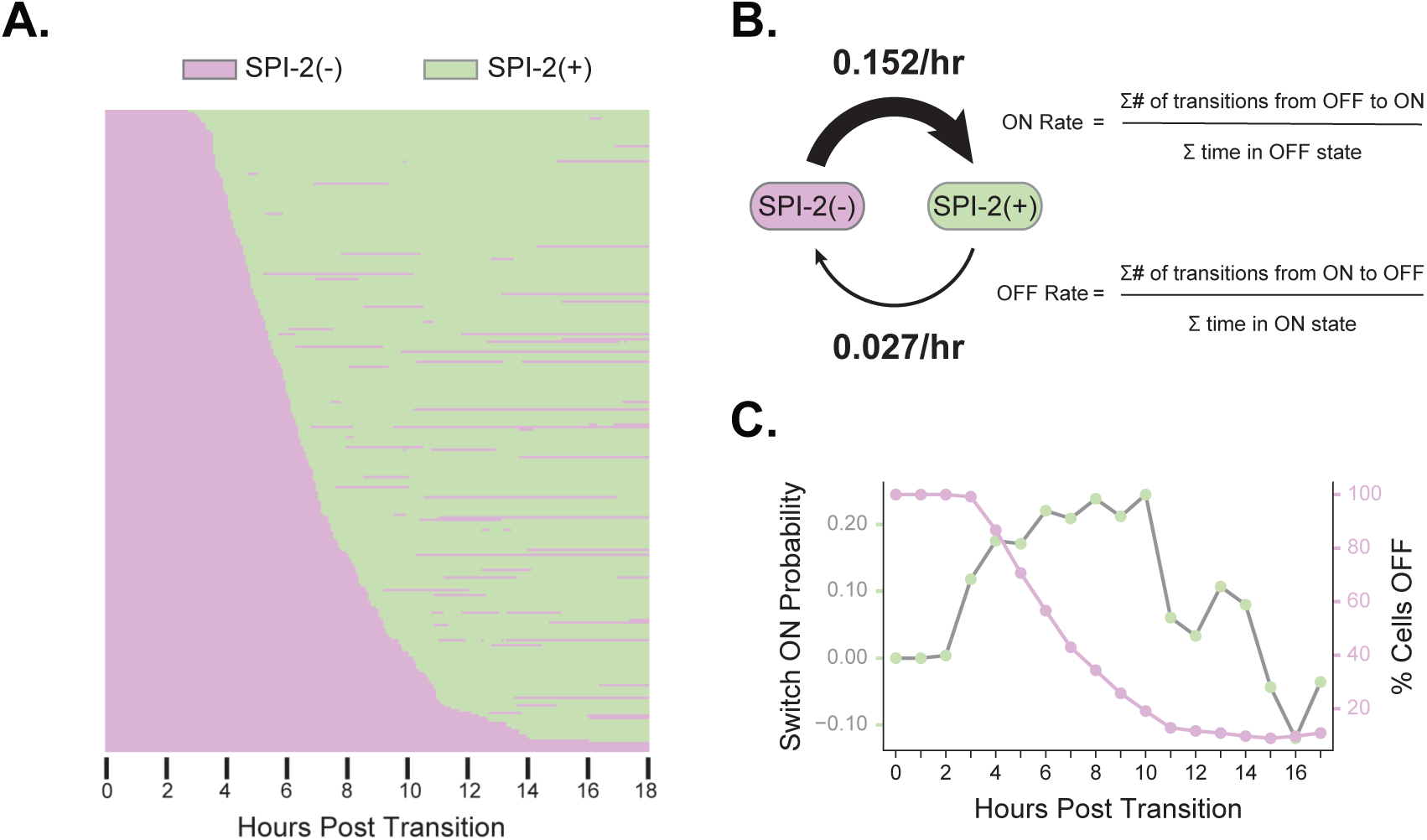
3. SPI-2 reporter activation is committed and equally probable across a range of timepoints (A) Mother cell data from Figure 6D was converted into a binary heatmap showing times in SPI-2(+) and SPI-2(-) states based on GFP(+) threshold as defined in Figure 6B. (B) Data from cells in (A) was used to calculate the population level SPI-2 ON and OFF switching rates based on time in each state. (C) Probability of SPI-2 transition into the ON state (shown in green) calculated on a per hour basis relative to the number of OFF cells at the start of each hour (shown in pink). Probability is equal to the difference in number of ON cells at the start and end of the hour divided by the number of cells OFF at the start of the hour. SPI-2 ON state transitions are predominantly committed and have a wide range of times (∼4-10 hours post transition into MgM-MES) during which they occur at similar probabilities.

**Figure S6.**
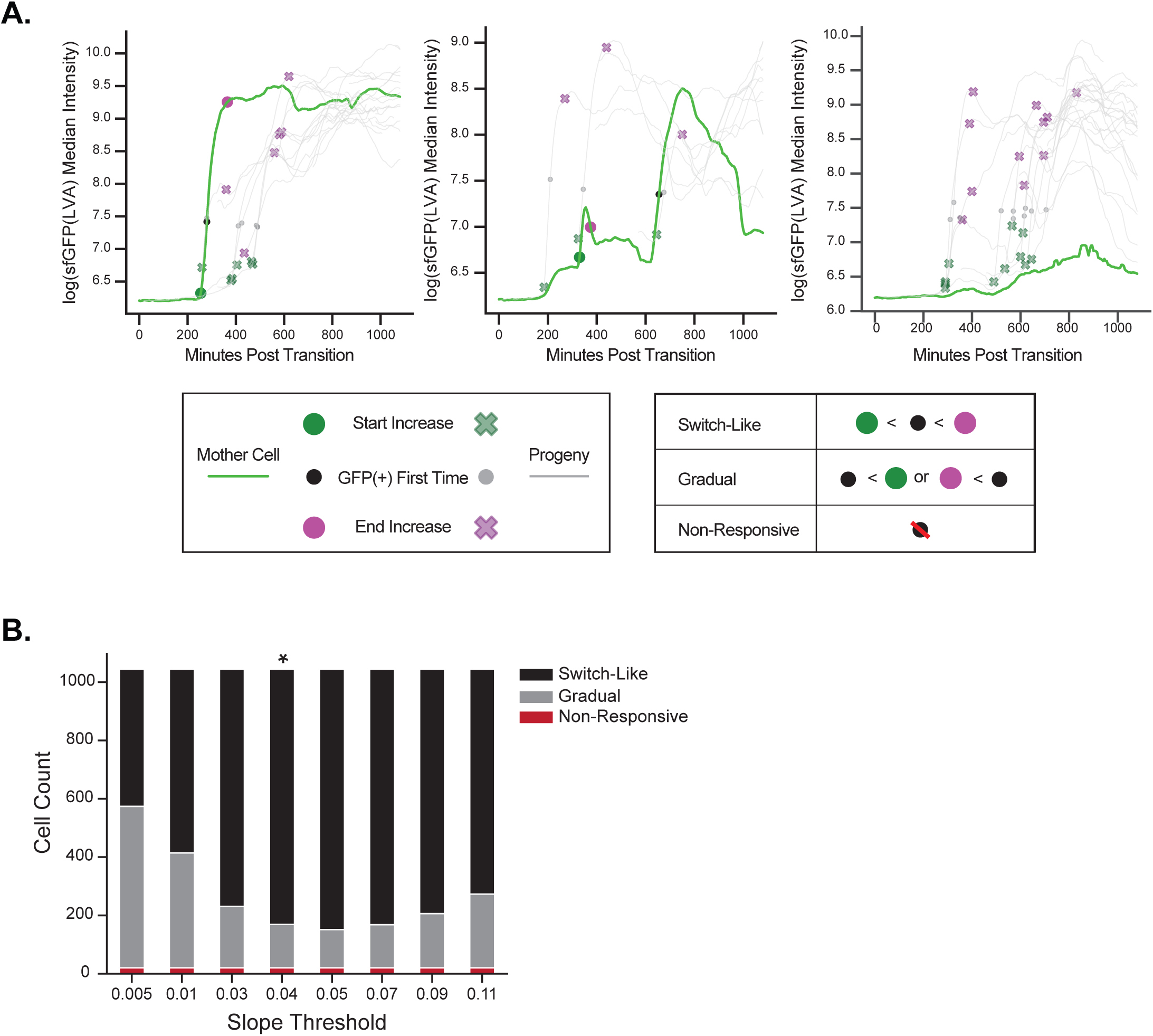
4. Detection and distribution of SPI-2 switching events based on sfGFP intensity slopes (A) SPI-2 reporter increase start times were defined based on the time at which the slope of the log(sfGFP(LVA) median intensity) increased above a threshold (0.04, see panel B). Reporter end increase times were defined as the point at which the slope dropped below the same threshold for at least 5 timepoints. Sample plots for representative mothers and all progeny cells with detected start and end increase times shown for each cell (Box 1). Cells were categorized on the basis of when the GFP(+) threshold was crossed relative to the start and end of the detected increase (Box 2). Switch-like cells were defined as cells in which the GFP(+) threshold was crossed between the start and end of the detected increase in GFP intensity. Gradual cells were defined as cells that crossed the GFP(+) threshold outside of the time window between the detected start and end increase. Non-responsive cells were defined as cells that never crossed the GFP(+) threshold. Mother cells (green) are representative of each of the three categories (from left to right, switch-like, gradual, and non-responsive). Cells that were born at intensities above the GFP(+) threshold or were tracked for fewer than 6 hours were not categorized. (B) Multiple slope thresholds were evaluated for their capacity to accurately detect switches and categorize cells into one of the three response types. The fraction of cells in each category was similar for thresholds between 0.03-0.07 and a threshold of 0.04 (asterisk, representative results in (A)) was selected as it most accurately detected major switching events and was used for all subsequent response type analysis.

**Figure S6.**
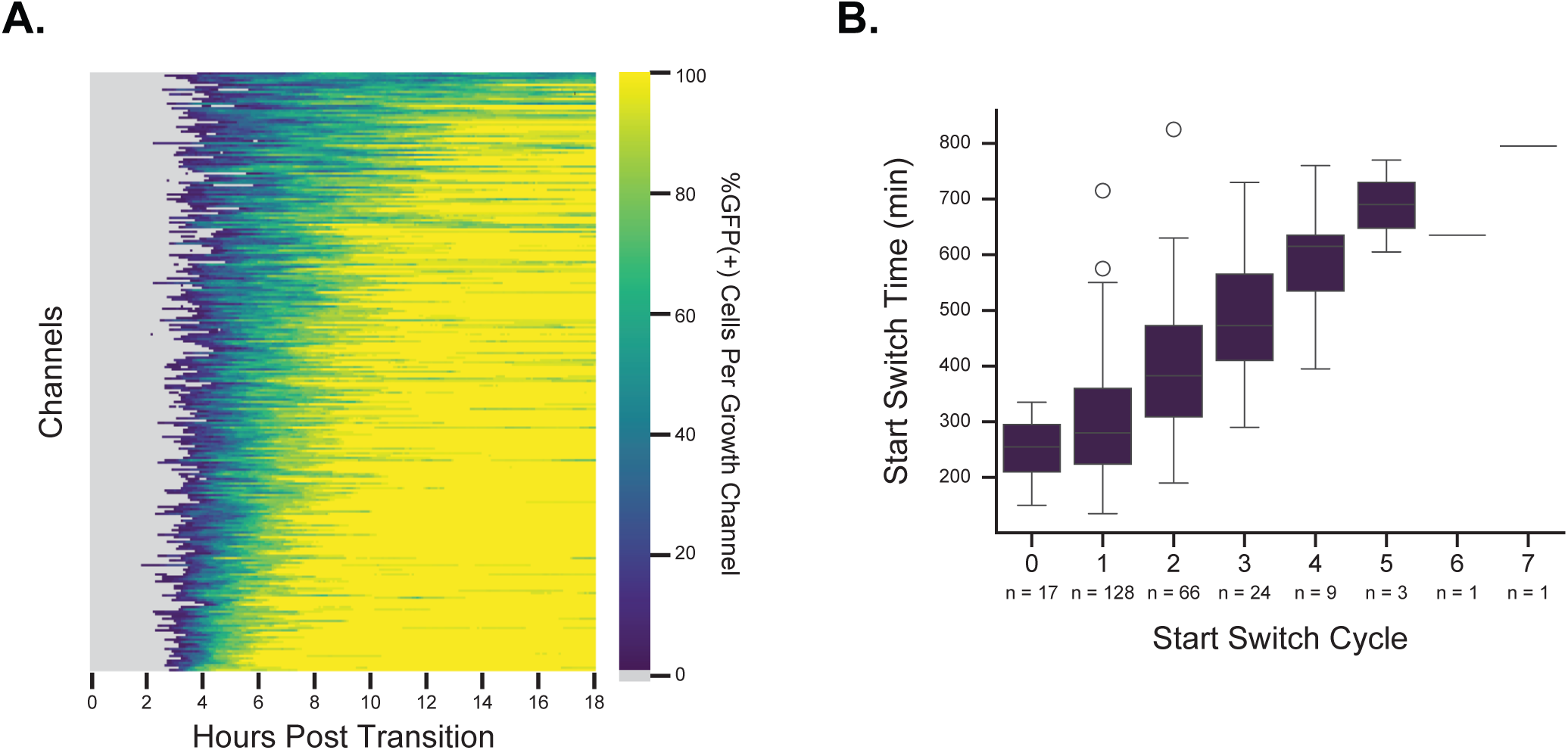
5. Single-cell response timing is not dependent on pre-existing variation or cell cycles number (A) Percent of cells within individual channels classified as SPI-2(+) based on the GFP(+) threshold are displayed as a heatmap across time. Early cell responses within a channel do not strongly correlate to the rate at which the rest of the channel becomes SPI-2(+). (B) The cycle number in which switch-like mother cells initiate SPI-2 activation as defined by the slope threshold (Figure S6.4). Responses are not linked to the number of cycles following the media transition.

**Figure S6.**
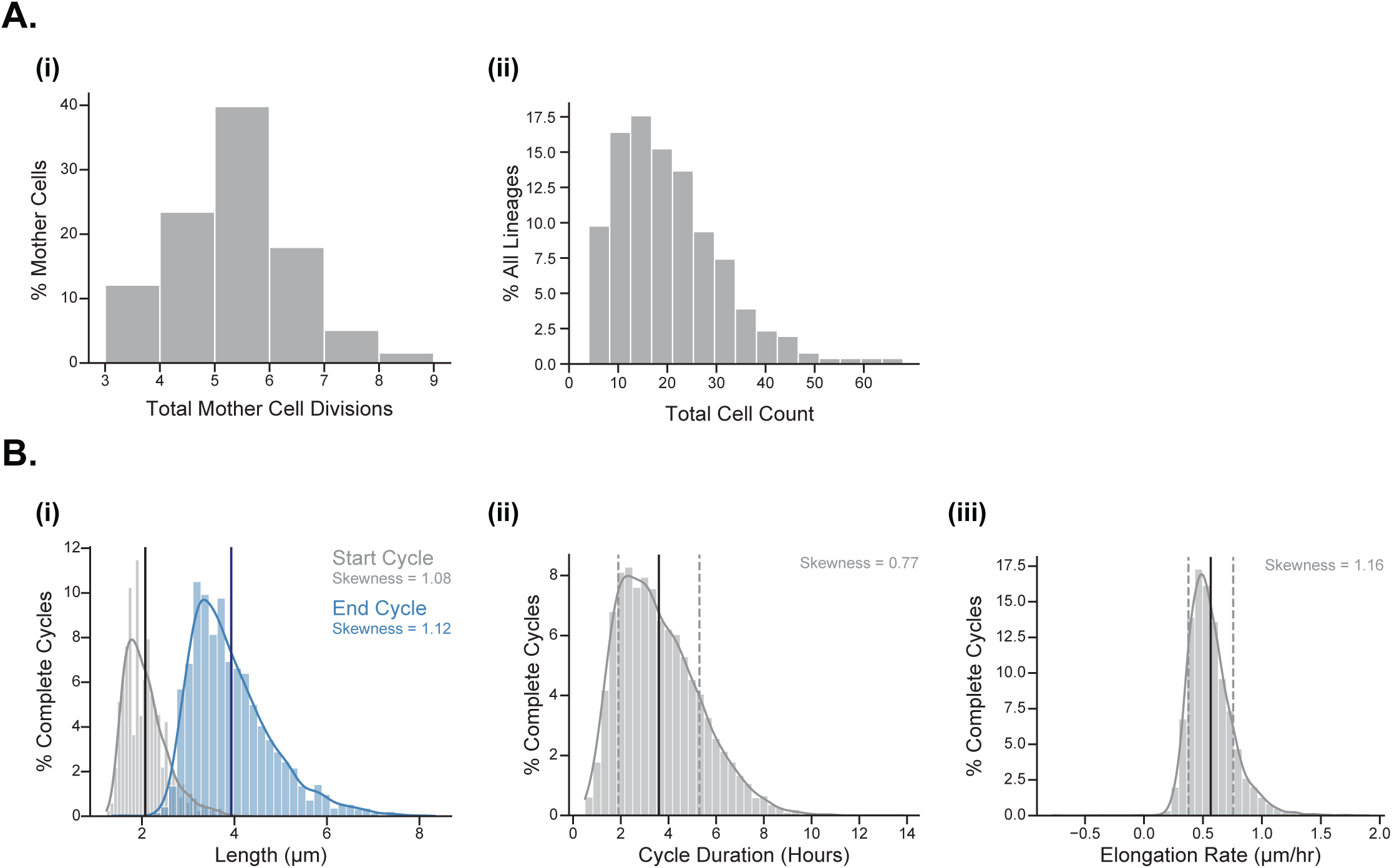
6. Features of cellular growth in SPI-2 inducing media in the DIMM (A) (i) The number of divisions mother cells experience and (ii) the total number of progeny tracked from each mother cell over 18 hours of growth in SPI-2 inducing media are shown, n=256 mother cells. Progeny count is limited by the length of the channel, as progeny are washed out and cannot be tracked through subsequent divisions in channels with rapidly dividing cells.(B) Distributions of features of complete cell cycles for all tracked cells within the DIMM experiments.

**Figure S6.**
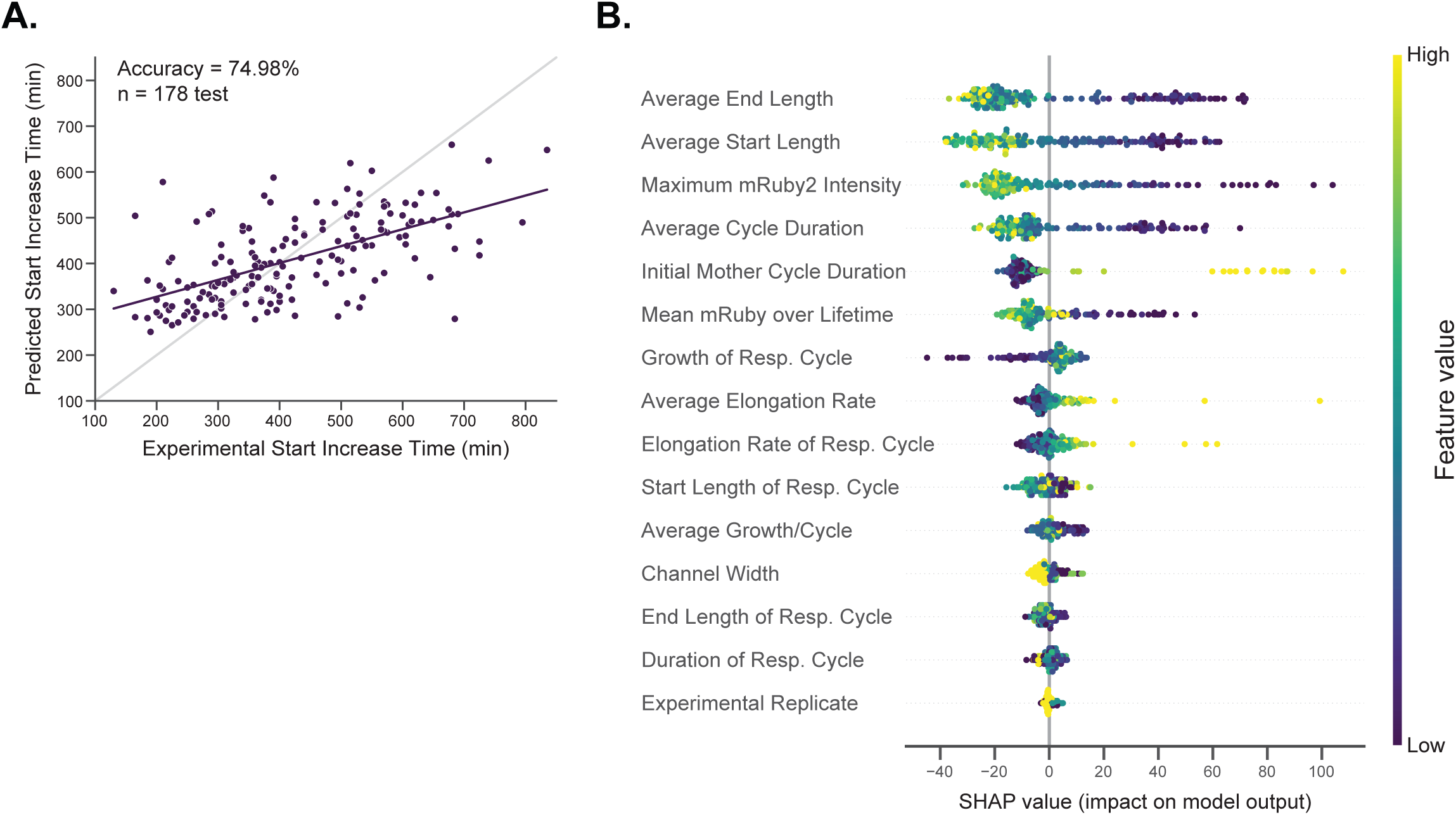
7. Cell growth features poorly predict SPI-2 activation time A Random Forest Regressor model was trained on 15 features of non-mother cells that were classified as switch-like and responded in a cycle that was fully capture by the experiment. (A) Experimental vs predicted start increase times from the model output. Accuracy metric is equal to 100 - MAPE value between experimental and predicted. Gray line represents y=x ideal prediction. Purple line is a linear fit of the data. n=178 cells. (B) SHAP values for all parameters used to train the model. Distance along the x-axis represents the contribution of the feature to the prediction, the color scale represents the value of the metric relative to all other values of the same metric, each dot is a predicted cell value.

**Figure S7.**
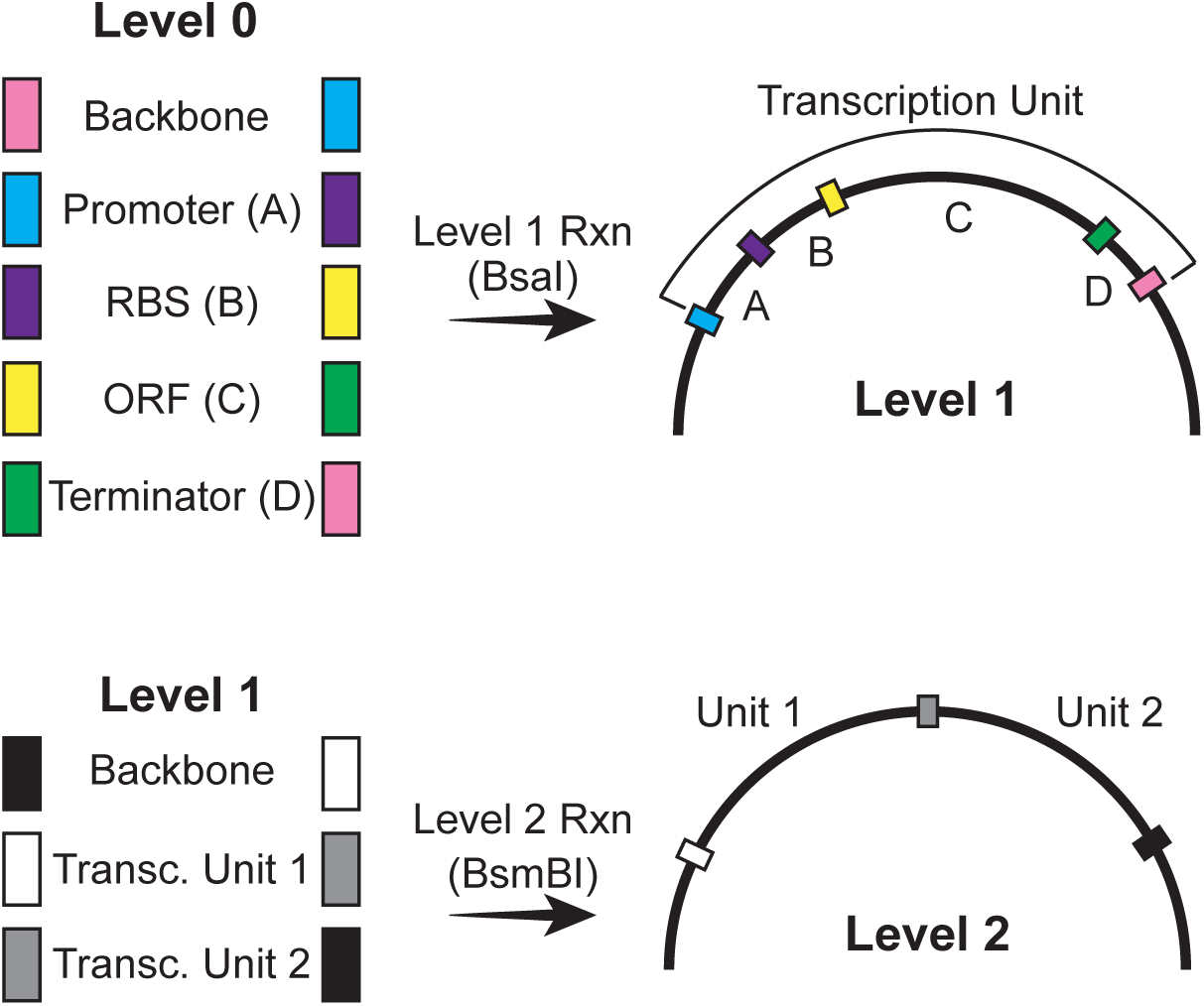
Golden-gate assembly of expression plasmids Plasmid assembly was based on the EcoFlex library. Level 0 plasmids containing transcription unit parts (promoter, RBS, ORF, terminator) are digested with BsaI and assembled via the compatible golden gate overhangs into a Level 1 expression plasmid. Level 1 plasmids are digested with BsmBI and assembled into a Level 2 plasmid containing two transcription units.

**Figure S8.**
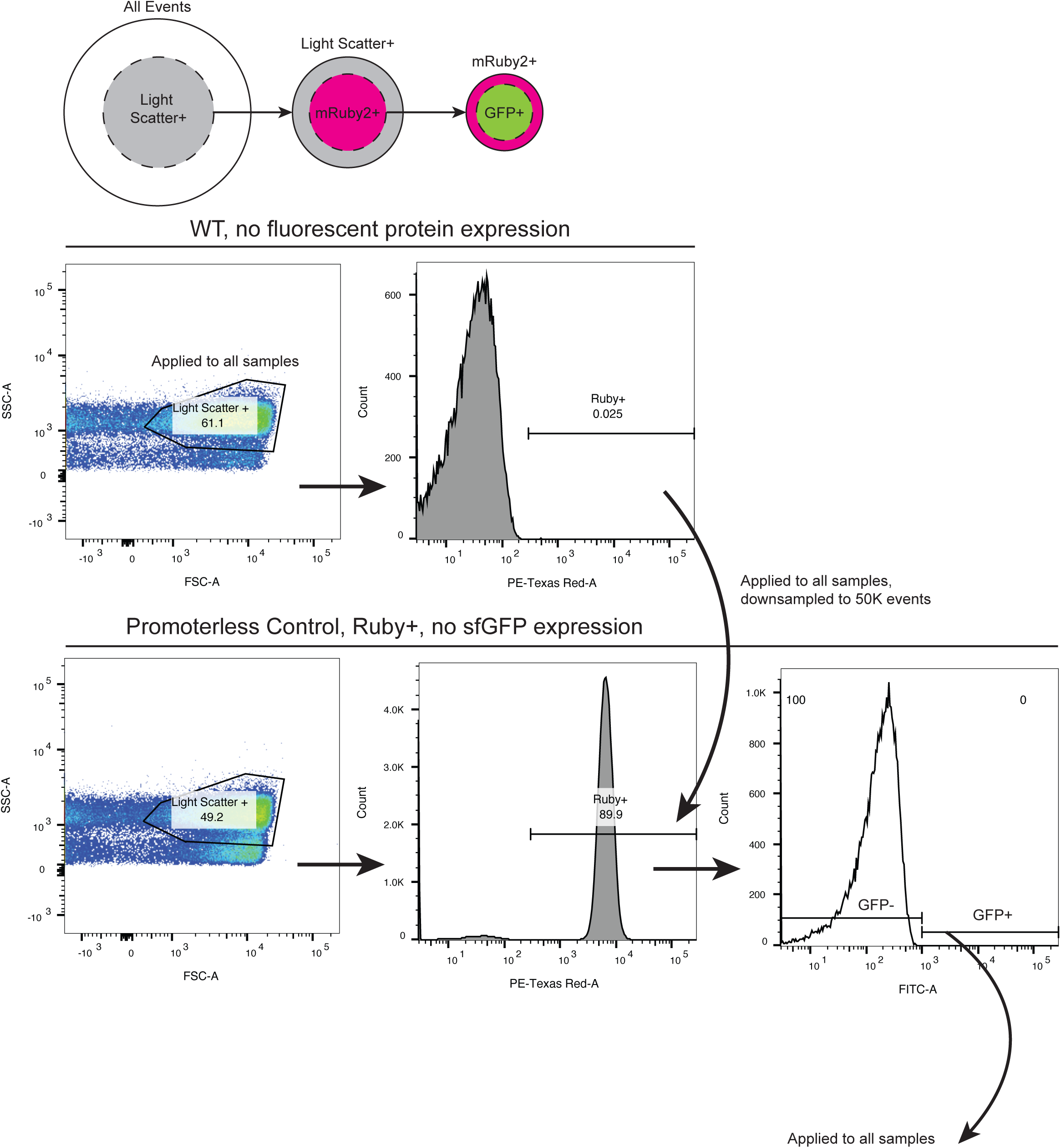
Flow cytometry gating methods Methods for gating and analyzing the SPI-2 reporters are shown. Top: Logic of the gates drawn. Middle: Control gates are drawn on non-fluorescent WT samples. The same light scatter gate is applied to all samples within the experiment. An mRuby2(+) gate is drawn on the WT populations such that less than 0.05 percent of the population is considered mRuby2(+). This gate is applied to all samples. All mRuby2(+) samples are DownSampled to 50,000 events. A GFP(+) gate is drawn on a GFP(-) control, and statistics are extracted from both populations.

## Notes

### Competing Interest Statement

The authors have declared no competing interest.

